# Soil Prokaryotic and Fungal Biome Structures Associated with Crop Disease Status across the Japan Archipelago

**DOI:** 10.1101/2023.08.31.555816

**Authors:** Hiroaki Fujita, Shigenobu Yoshida, Kenta Suzuki, Hirokazu Toju

## Abstract

Archaea, bacteria, and fungi in the soil are increasingly recognized as determinants of agricultural productivity and sustainability. A crucial step for exploring soil microbiomes with high ecosystem functions is to perform statistical analyses on potential relationship between microbiome structure and functions based on comparisons of hundreds or thousands of environmental samples collected across broad geographic ranges. In this study, we integrated agricultural field metadata with microbial community analyses by targeting > 2,000 soil samples collected along a latitudinal gradient from cool-temperate to subtropical regions in Japan (26.1– 42.8 °N). The data involving 632 archaeal, 26,868 bacterial, and 4,889 fungal operational taxonomic units detected across the fields of 19 crop plant species allowed us to conduct statistical analyses (permutational analyses of variance, generalized linear mixed models, and randomization analyses) on relationship among edaphic factors, microbiome compositions, and crop disease prevalence. We then examined whether the diverse microbes form species sets varying in potential ecological impacts on crop plants. A network analysis suggested that the observed prokaryotes and fungi were actually classified into several species sets (network modules), which differed substantially in associations with crop disease prevalence. Within the network of microbe-to-microbe coexistence, ecologically diverse microbes, such as an ammonium-oxidizing archaeum, an antibiotics-producing bacterium, and a potentially mycoparasitic fungus, were inferred to play key roles in shifts between crop-disease-promotive and crop-disease-suppressive states of soil microbiomes. The bird’s-eye view of soil microbiome structure will provide a basis for designing and managing agroecosystems with high disease-suppressive functions.

**IMPORTANCE:** Understanding how microbiome structure and functions are organized in soil ecosystems is one of the major challenges in both basic ecology and applied microbiology. Given the ongoing worldwide degradation of agroecosystems, building frameworks for exploring structural diversity and functional profiles of soil microbiomes is an essential task. Our study provides an overview of cropland microbiome states in light of potential crop-disease-suppressive functions. The large dataset allowed us to explore highly functional species sets that may be stably managed in agroecosystems. Furthermore, an analysis of network architecture highlighted species that are potentially used to cause shifts from disease-prevalent states of agroecosystems to disease-suppressive states. By extending the approach of comparative analyses towards broader geographic ranges and diverse agricultural practices, agroecosystem with maximized biological functions will be further explored.

---

The ongoing global-scale degradation of agroecosystems is threatening food production (1, 2). Maximizing the functions of microbial communities (microbiomes) is a prerequisite for building bases of sustainable agriculture (3–7). Archaea, bacteria, and fungi in the soil drive cycles of carbon, nitrogen, and phosphorus within agroecosystems (8–12). Many of those microbes also work to promote crop plant’s tolerance to drought and high temperature stresses as well as resistance to pests and pathogens (13–18). Importantly, those microbes vary greatly in their physiological impacts on crop plants (19–21). Therefore, gaining insights into soil microbiome compositions is an essential starting point for managing resource-use efficient and disease-tolerant agroecosystems.

Since the emergence of high-throughput DNA sequencing, a number of studies have revealed taxonomic compositions of prokaryotes and/or fungi in agroecosystem soil (22–24). Those studies have explored microbial species that potentially support crop plant growth and/or prevent crop plant disease (9, 16, 25, 26). Meanwhile, each of the previous studies has tended to focus on specific crop plant species in specific farm fields (27), although there are some exceptionally comprehensive studies comparing multiple research sites (15, 22). Therefore, generality in relationship between microbiome structure and functions remain to be examined in broader contexts [cf. global-scale analyses of soil microbiomes in natural ecosystems (28–31)]. In other words, we still have limited knowledge of general patterns and features common to soil microbiomes with high crop yield or those with least crop disease risk. Thus, statistical analyses comparing microbiome structure among diverse crop plants across broad geographic ranges (15, 22) are expected to deepen our understanding of microbial functions in agroecosystems. In particular, comparative studies of thousands of soil samples covering a wide range of latitudes will provide opportunities for finding general properties common to microbial communities with plant-growth-promoting or crop-disease-suppressive functions across diverse climatic conditions.

Large datasets of soil microbiomes will also allow us to estimate interspecific interactions between microbial species (3, 32, 33). Archaea, bacteria, and fungi in soil ecosystems potentially form entangled webs of facilitative or competitive interactions, collectively determining ecosystem-level functions such as the efficiency of nutrient cycles and the prevalence of plant pathogens (34, 35). In fact, ecological network studies have inferred how sets of microbial species could respond to the outbreaks or experimental introductions of crop plant pathogens (36–38). Although various statistical platforms for deciphering the architecture of such microbial interaction networks have been proposed (32, 39), hundreds or more of microbial community samples are required to gain reliable inferences on interactions that reproducibly occur in real ecosystems (40). Thus, datasets consisting of thousands of soil samples collected across a number of local ecosystems will provide fundamental insights into how soil ecological processes are driven by cross-kingdom interactions involving archaea, bacteria, and fungi.

In this study, we conducted a comparative analysis of agroecosystem soil microbiomes based on > 2,000 soil samples collected from subtropical to cool-temperate regions across the Japan Archipelago. Based on the amplicon sequencing dataset representing farm fields of 19 crop plant species, we profiled prokaryotic and fungal community compositions in conventional agricultural fields in Japan. By compiling the metadata of the soil samples, we then examined biotic and abiotic factors explaining diversity in the prevalence of crop disease. The soil microbiome dataset was further used to infer the structure of a microbe-to-microbe coexistence network consisting of diverse archaea, bacteria, and fungi. Specifically, we examined whether the network architecture was partitioned into compartments (modules) of closely interacting microbial species. In addition, we tested the hypothesis that such network modules could differ in their positive/negative associations with crop plant disease/health status. To explore prokaryotic and fungal species keys to manage agroecosystem structure and functions, we further explored “core” or “hub” species that were placed at the central positions within the inferred microbial interaction network. Overall, this study provides an overview of soil microbial diversity of cropland soil across a latitudinal gradient, setting a basis for diagnosing soil ecosystem status and identifying sets of microbes to be controlled in sustainable crop production.

## RESULTS

### Diversity of agroecosystem microbiomes

We compiled the field metadata of 2,903 soil samples collected in the research projects of National Agricultural and Food Research Organization (NARO), Japan. The bulk soil of farmlands was sampled from subtropical to cool-temperate regions (26.1–42.8 °N) across the Japan Archipelago from 2006 to 2014, targeting 19 crop plant species (Fig. 1A; Data S1). Most of the croplands were managed with conventional agricultural practices (characterized by intensive tillage and chemical fertilizer/pesticide application), while some were experimentally controlled as organic agricultural fields. The metadata (Data S1) included the information of chemical [e.g., pH, electrical conductivity, carbon/nitrogen (C/N) ratio, and available phosphorous concentration], physical (e.g., soil taxonomy), and biological (e.g., crop disease level) properties, providing a platform for profiling ecosystem states of cropland soil.

**Fig. 1.**
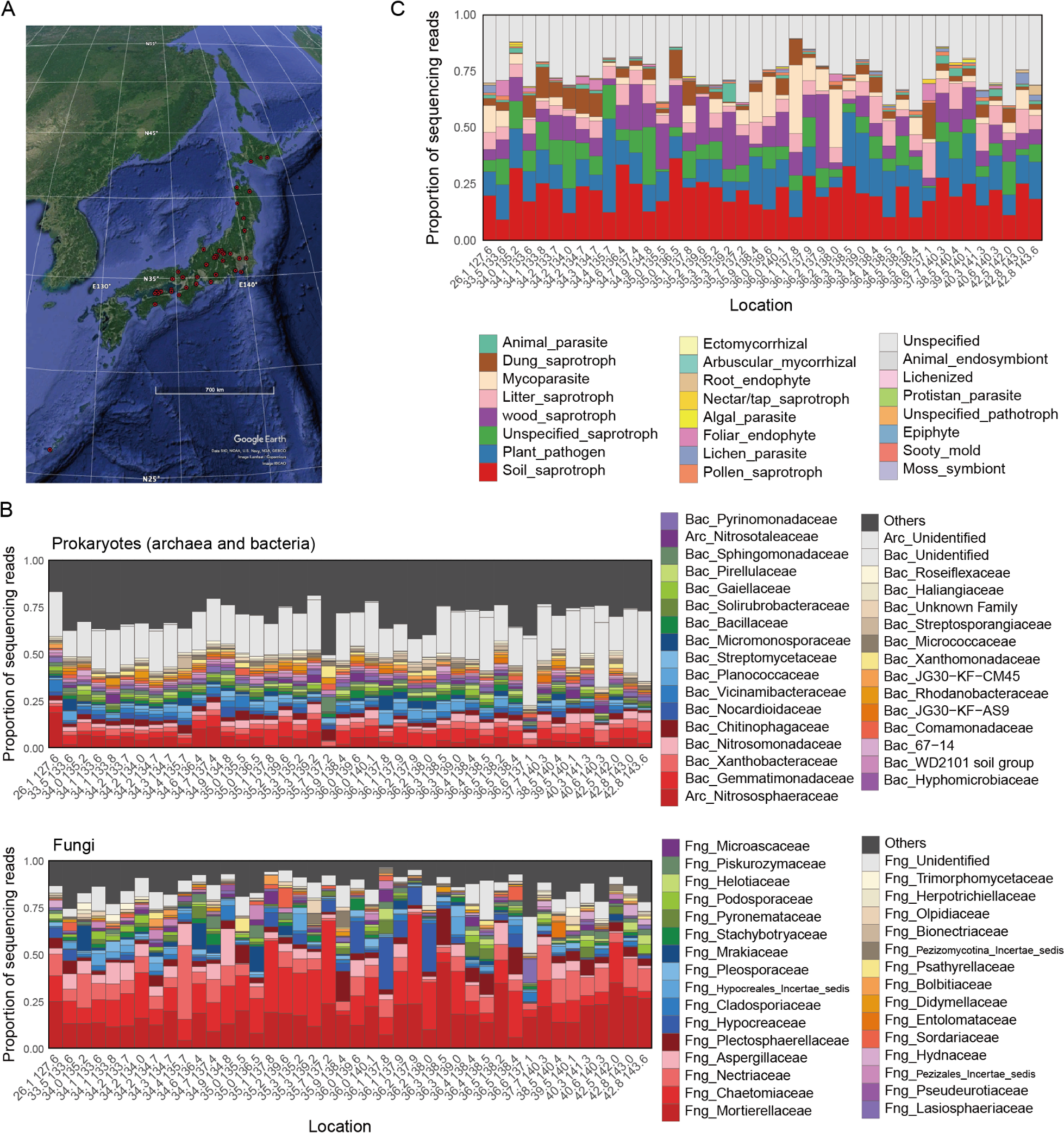
Comparison of soil microbiome structure across the Japan Archipelago. (A) Map of research sites across the Japan Archipelago. The 2,903 soil samples were grouped into 42 research sits were when their latitude and longitude profiles were round to one decimal place. (B) Taxonomic compositions of prokaryotes (archaea and bacteria; top) and fungi (bottom) at the family level. See Fig. S1 for results at the genus, order, and class levels. (C) Compositions of functional groups of fungi.

To integrate the metadata with the information of microbial community structure, we performed DNA metabarcoding analyses of both prokaryotes (archaea and bacteria) and fungi. After a series of quality filtering, prokaryotic and fungal community data were obtained from 2,676 and 2,477 samples, respectively. In total, 632 archaeal operational taxonomic units (OTUs) representing 22 genera (24 families), 26,868 bacterial OTUs representing 1,120 genera (447 families), and 4,889 fungal OTUs representing 1,190 genera (495 families) were detected (Fig. 1B; Fig. S1).

The prokaryotic communities lacked apparently dominant taxa at the genus and family levels (Fig. 1B). In contrast, the fungal communities were dominated by fungi in the families Mortierellaceae, Chetomiaceae, and Nectriaceae, depending on localities (Fig. 1B). A reference database profiling of fungal functional groups suggested that the fungal communities were dominated by soil saprotrophic and plant pathogenic fungi (Fig. 1C) as characterized by the dominance of *Mortierella* and *Fusarium* at the genus level (Fig. S1). Meanwhile, mycoparasitic fungi had exceptionally high proportions at some research sites, as represented by the dominance of *Trichoderma* (Hypocreaceae) at those sites (Fig. 1B; Fig. S1).

### Microbiome structure and crop disease prevalence

Compiling the metadata of edaphic factors, we found that variation in the community structure of prokaryotes and fungi was significantly explained by crop plant identity and soil taxonomy as well as by soil chemical properties such as pH, electrical conductivity, and C/N ratio, (Fig. 2A-B; Figs. S2-3; Table 1). In addition, the ratio of prokaryotic abundance to fungal abundance (see Materials and Methods for details) was associated with both prokaryotic and fungal community structure (Table 1). Nonetheless, the explanatory powers of these variable were all small as indicated by the low *R*^2^ values (Table 1).

**Fig. 2.**
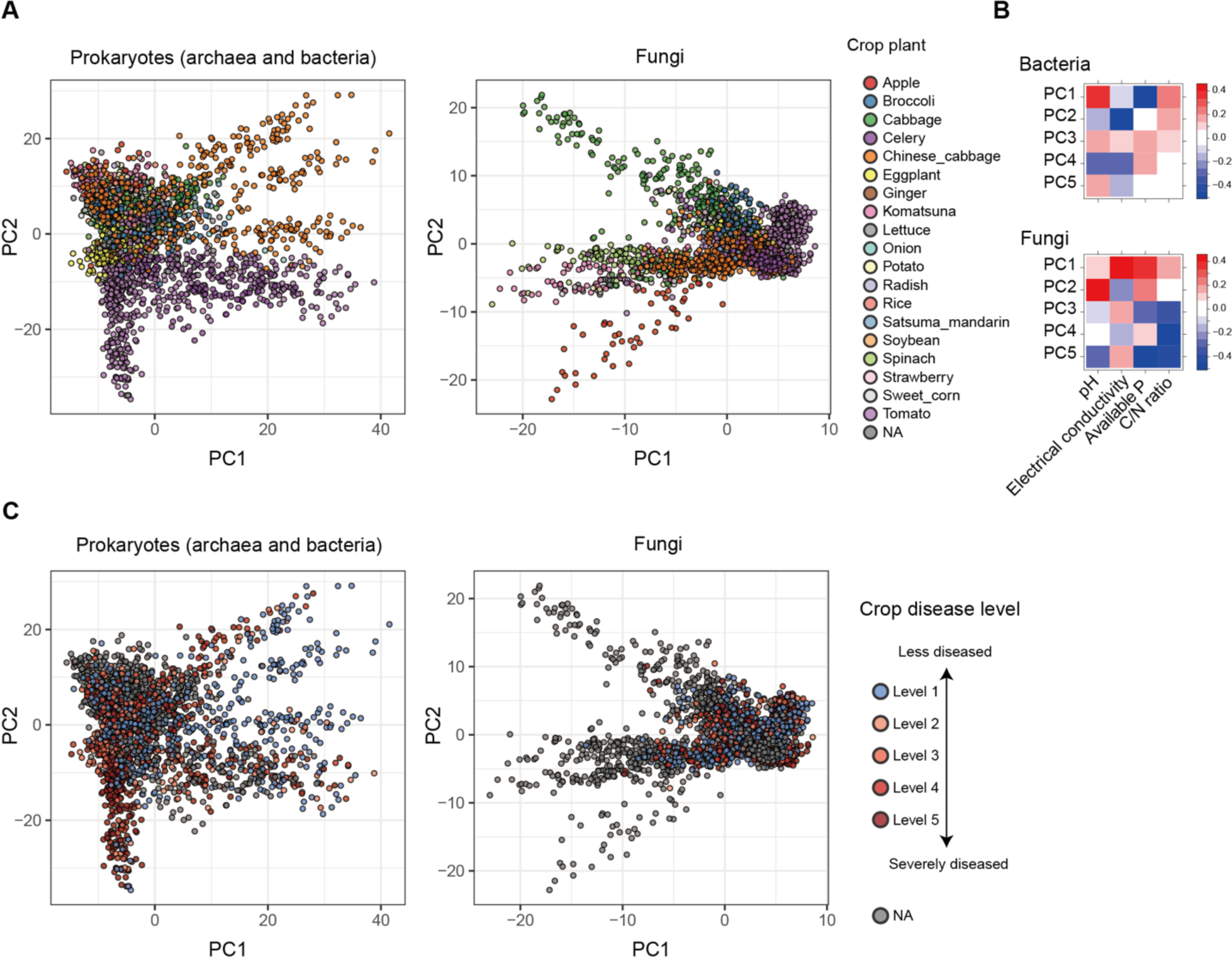
Dimensions of soil microbiome structure. (A) Prokaryote and fungal community structure. Principal co-ordinate analyses (PCA) were performed based on OTU-level compositional matrices respectively for the prokaryotic and fungal communities. The identify of crop plants is shown by colors. See Figures S2-3 for relationship between community structure and environmental factors. (B) Correlation between PCA scores and soil environmental factors. (C) Crop disease level and microbial community structure. On the PCA surface, crop disease level (see Materials and Methods) is indicated.

**Table 1.** Effects of environmental variables on prokaryotic/fungal community structure. For each set of categorical/continuous environmental variables, a PERMANOVA was performed for each of the prokaryotic and fungal community datasets.

| Model | Dataset | Variable | df | $R^2$ | $F$ | $P$ |
| --- | --- | --- | --- | --- | --- | --- |
| Categorical variables | Prokaryotes | Research site | 34 | 0.051 | 5.13 | < 0.001 |
|  |  | Month | 11 | 0.014 | 4.36 | < 0.001 |
|  |  | Crop | 16 | 0.006 | 1.21 | 0.030 |
|  |  | Former crop | 22 | 0.024 | 3.78 | < 0.001 |
|  |  | Soil category | 10 | 0.006 | 2.12 | < 0.001 |
|  |  | Residual | 2301 | 0.673 |  |  |
|  |  | Total | 2400 | 1.000 |  |  |
|  | Fungi | Research site | 33 | 0.064 | 6.91 | < 0.001 |
|  |  | Month | 11 | 0.012 | 3.96 | < 0.001 |
|  |  | Crop | 15 | 0.006 | 1.54 | < 0.001 |
|  |  | Former crop | 20 | 0.023 | 4.14 | < 0.001 |
|  |  | Soil category | 10 | 0.005 | 1.75 | < 0.001 |
|  |  | Residual | 2109 | 0.591 |  |  |
|  |  | Total | 2206 | 1.000 |  |  |
| Continuous variables | Prokaryotes | pH | 1 | 0.011 | 16.52 | < 0.001 |
|  |  | Electrical conductivity | 1 | 0.009 | 14.09 | < 0.001 |
|  |  | Available P | 1 | 0.009 | 13.81 | 0.626 |
|  |  | C/N ratio | 1 | 0.004 | 6.19 | < 0.001 |
|  |  | Latitude | 1 | 0.006 | 9.00 | < 0.001 |
|  |  | Longitude | 1 | 0.008 | 11.41 | < 0.001 |
|  |  | Prokaryote/fungus ratio | 1 | 0.004 | 5.98 | < 0.001 |
|  |  | Residual | 1408 | 0.936 |  |  |
|  |  | Total | 1415 | 1.000 |  |  |
|  | Fungi | pH | 1 | 0.013 | 19.64 | < 0.001 |
|  |  | Electrical conductivity | 1 | 0.011 | 17.49 | < 0.001 |
|  |  | Available P | 1 | 0.009 | 13.38 | 0.477 |
|  |  | C/N ratio | 1 | 0.008 | 12.74 | < 0.001 |
|  |  | Latitude | 1 | 0.016 | 25.58 | < 0.001 |
|  |  | Longitude | 1 | 0.017 | 26.37 | 0.230 |
|  |  | Prokaryote/fungus ratio | 1 | 0.009 | 13.61 | < 0.001 |
|  |  | Residual | 1408 | 0.904 |  |  |
|  |  | Total | 1415 | 1 |  |  |

Both prokaryotic and fungal community structure was significantly associated with the severity of crop disease (Fig. 2C; Table 2). Specifically, the crop plants’ disease/health status (disease level 1 vs. disease levels 2-5; see Materials and Methods) was explained by some of the principal components (PCs) defined based on prokaryotic/fungal community structure (Fig. 2B).

**Table 2.** Relationship between prokaryotic/fungal community structure on the disease level of crop plants. A GLMM of crop plants’ disease level (disease level 1 vs. disease levels 2-5) with a logit-link function and binomial errors was constructed by setting principal components of prokaryotic/fungal community structure (Fig. 2) as explanatory variables (fixed effects). The identity of experimental/research purposes, sampling month, and crop plant species were included as random effects in the GLMM.

| Dataset | Variable | *z* | *P* |
| --- | --- | --- | --- |
| Prokaryotes<br>(*N* = 1,379) | PC1 | 1.59 | 0.1111 |
|  | PC2 | -1.65 | 0.1000 |
|  | PC3 | 1.82 | 0.0684 |
|  | PC4 | -2.32 | 0.0205 |
|  | PC5 | 3.98 | 0.0001 |
| Fungi<br>(*N* = 1,320) | PC1 | 1.52 | 0.1281 |
|  | PC2 | 1.39 | 0.1656 |
|  | PC3 | -2.11 | 0.0348 |
|  | PC4 | 2.62 | 0.0089 |
|  | PC5 | -0.84 | 0.4002 |

### Microbes associated with crop disease/health status

We explored microbial OTUs whose prevalence are associated with crop plant disease/health status. Based on a randomization analysis, prokaryotic/fungal OTUs whose distribution is biased in samples representing the minimal crop disease level (disease level 1) were screened (Fig. S4).

To examine whether the OTUs highlighted in the across-Japan spatial scale could actually show tight associations with crop disease status at local scales, the randomization analysis was performed as well in each of the six sub-datasets representing unique combinations of research sites, crop plant species, and experimental/research purposes (Data S2). Statistically significant specificity for crop disease level (FDR < 0.025; two-tailed test) was observed for at least one OTU in five of the six sub-datasets (Data S2). Among them, exceptionally strong specificity to the minimal crop disease level (standardized specificity score ≥ 6.0; FDR < 0.0001) was detected in two sub-datasets (Table 3). The relative abundance of these OTUs tightly associated with crop disease level across samples within each sub-dataset (Fig. 3).

**Fig. 3.**
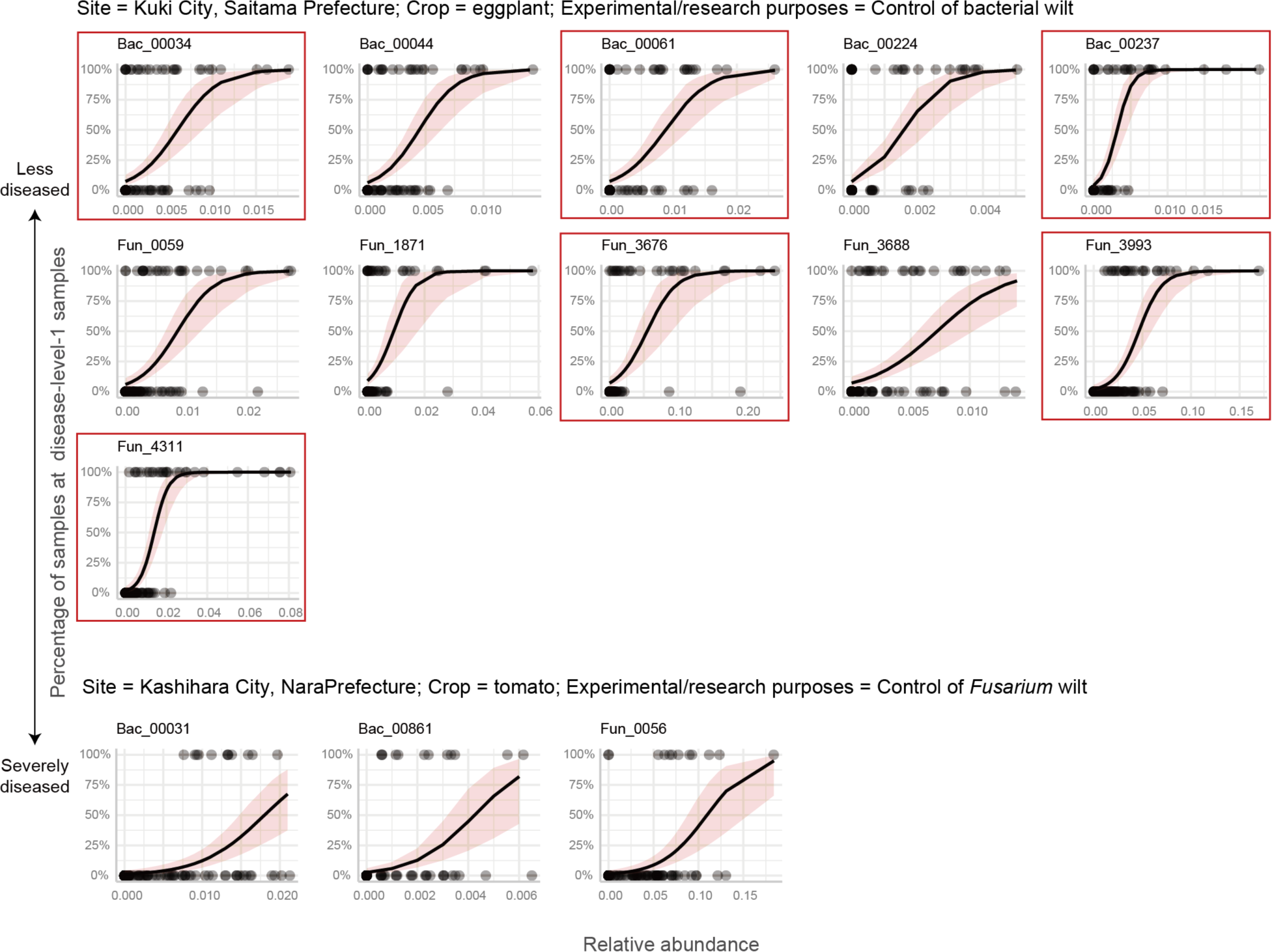
Relationship between OTU abundance and crop plant health. Among the six sub-datasets representing unique crop plant × site combinations, OTUs showing strongest specificity to the minimal crop disease level (*z*-standardized specificity to disease level 1 ≥ 6.0) were observed in two sub-datasets (“eggplant in Kuki City” and “tomato in Kashihara City”; Table 4; see Data S2 for full results). For each OTU in each sub-dataset, generalized linear model with a logit function and binomial errors was constructed to examine relationship between OTU relative abundance and crop disease level (level 1 vs. levels 2-5). All the regression lines are statistically significant (FDR < 0.0001). The OTUs exhibiting statistically significant specificity to disease level 1 in the analysis with the entire dataset (FDR < 0.025; two-tailed test; Fig. S4-5) are highlighted with red squares.

**Table 3.**
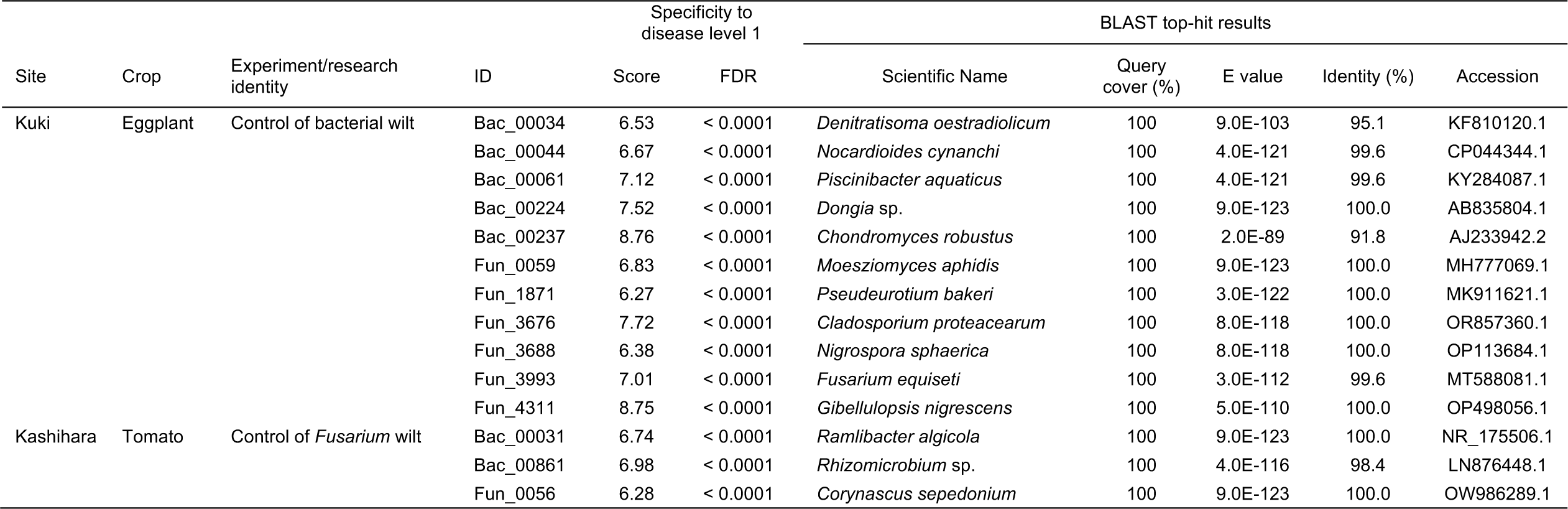
Prokaryotic and fungal OTUs showing highest associations with crop health status within local croplands. Among the six sub-datasets representing unique combinations of research sites, crop plant species, and research experimental/research purposes, OTUs showing strongest specificity to the minimal crop disease level (*z*-standardized specificity to disease level 1 ≥ 6.0) were observed in two sub-datasets (“eggplant in Kuki City” and “tomato in Kashihara City”). The OTUs are shown with the NCBI BLAST top-hit results. See Data S2 for the full results.

**Table 4.**
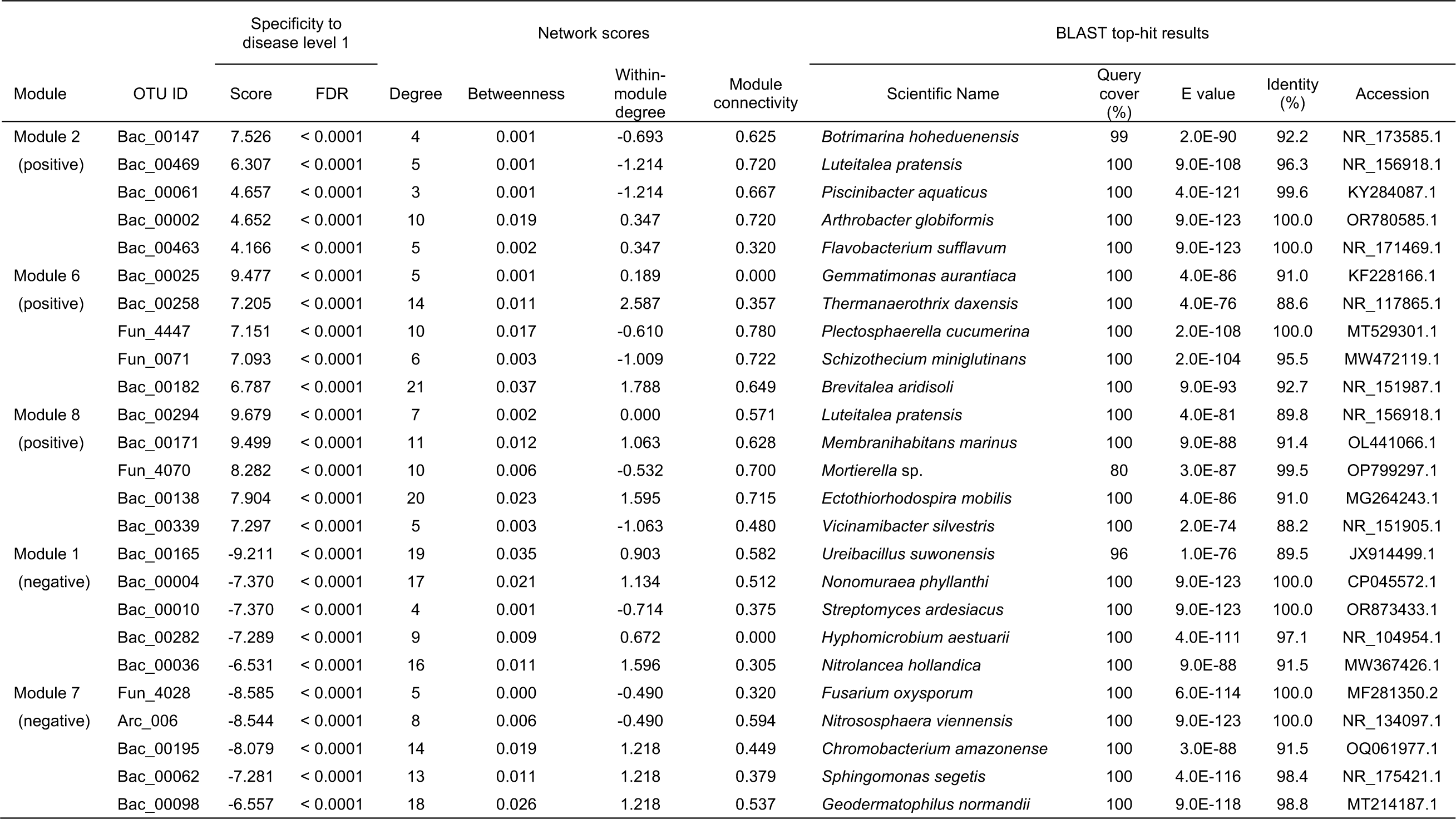
Representative prokaryotic and fungal OTUs in network modules with highly positive/negative associations with crop plant health. In each of the modules 2, 6, and 8 (Fig. 4), the top-five OTUs with the highest specificity to the minimal crop disease level (specificity to disease level 1; see Fig. S4 for the relationship between the specificity score and FDR). For each OTU, network degree, betweenness centrality, within-module degree (*z*-standardized), and among-module connectivity (Fig. 6) are presented with the NCBI BLAST top-hit results. Likewise, in each of the modules 1 and 7 (Fig. 4), the top-five OTUs negatively associated with the minimal crop disease level are shown. See Data S3 for the full results.

### Microbe-to-microbe network

We then examined the network architecture of potential microbe-to-microbe interactions within the soil microbiomes. The inferred network of coexistence was subdivided into several modules, in which archaeal, bacterial, and fungal OTUs sharing environmental preferences and/or those in positive interactions were linked with each other (Figs. 3A and S5-8). The network modules differed considerably in their associations with crop-plant disease level (Fig. 4B; Fig. S5; Data S3). Modules 2, 6, and 8, for example, were characterized by microbes associated with least disease level. Module 6, which showed the highest mean specificity to the minimal crop disease level (Fig. 4B), included a bacterium allied to the genus *Gemmatimonas* (Bac_00025), that allied to the genus *Thermanaerothrix* (Bac_00258), and a *Plectosphaerella* fungus (Fun_4447) (Table 4). In contrast to these modules, Modules 1 and 7 were constituted by microbes negatively associated with crop plant health (Fig. 4B). Module 1 included a bacterium distantly allied to the genus *Ureibacillus* (Bac_00165), a *Nonomuraea* bacterium (Bac_00004), and a *Streptomyces* bacterium (Bac_00010), while Module 7 involved a *Fusarium* fungus (Fun_4028) and a *Nitrososphaera* archaeum (Arc_006) (Table 4).

**Fig. 4.**
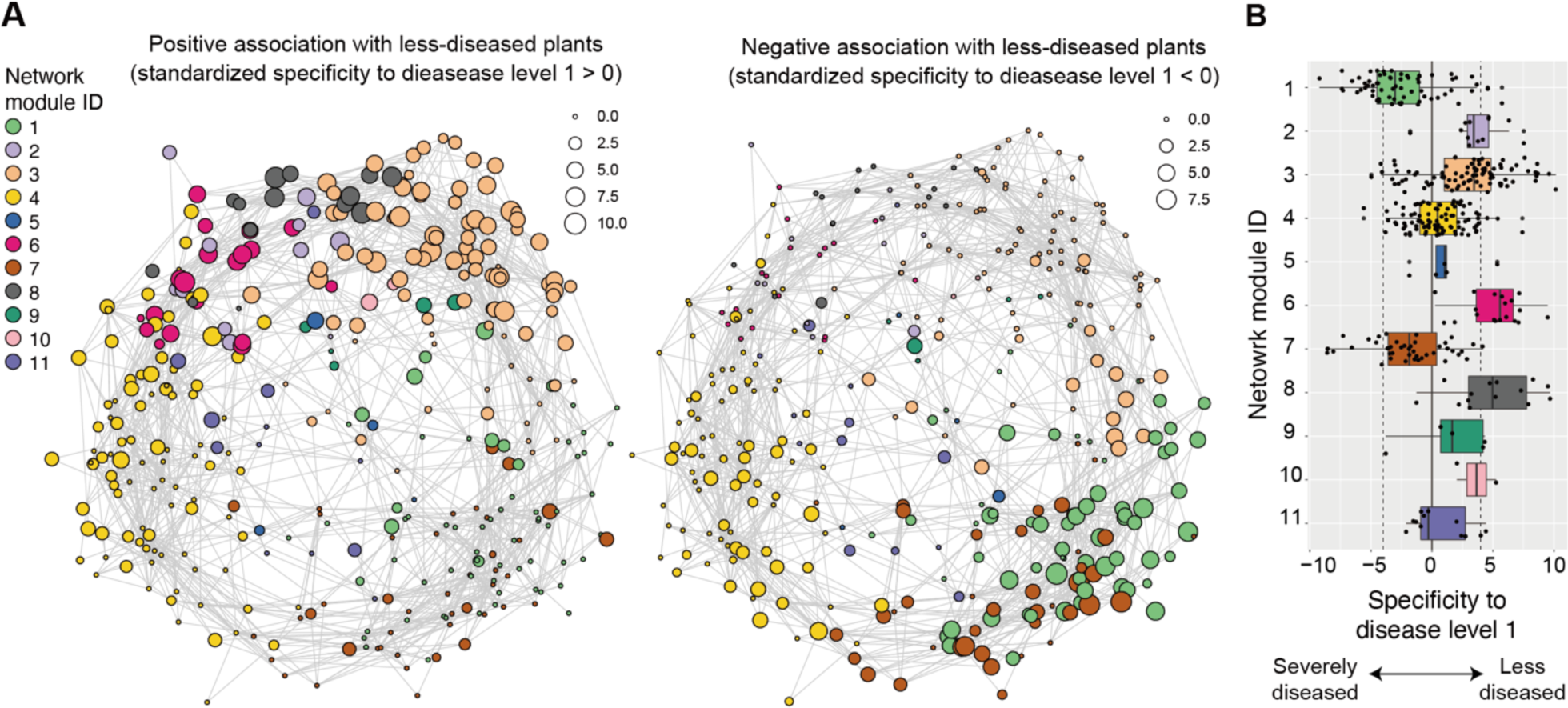
Architecture of microbe-to-microbe network. (A) Co-occurrence networks of archaea, bacteria, and fungi. Specificity of occurrences to disease-level-1 (the lowest disease level) samples (Figs. S4-5) is shown for each OTU within the network. The specificity is shown as node size separately for positive (left) and negative (right) associations with least-diseased states of crop plants. Colors indicate network modules, in which microbial OTUs in commensalistic/mutualistic interactions and/or those sharing environmental preferences are densely linked with each other. See Figure S8 for taxonomy (archaea, bacteria, or fungi) of respective nodes. (B) Characteristics of network modules. Mean specificity to the minimal crop disease level (disease level 1; left in the panel A) is shown for each network module.

### Core species within the microbial network

We next explored microbial OTUs that potentially have great impacts on community- or ecosystem-scale processes based on an analysis of the microbe-to-microbe network architecture (Data S3). Among the microbes disproportionately found from the samples with the minimal crop disease level, a Pyrinomonadaceae bacterium allied to the genus *Brevitalea* (Bac_00182 in Module 6; Table 4), for example, showed a high betweenness centrality score (Fig. 5). Meanwhile, among the microbes negatively associated with crop health status, a bacterium distantly allied to the genus *Ureibacillus* (Bac_00165 in Module 1; Table 4) was inferred to be located at a central position within the network (Fig. 5).

**Fig. 5.**
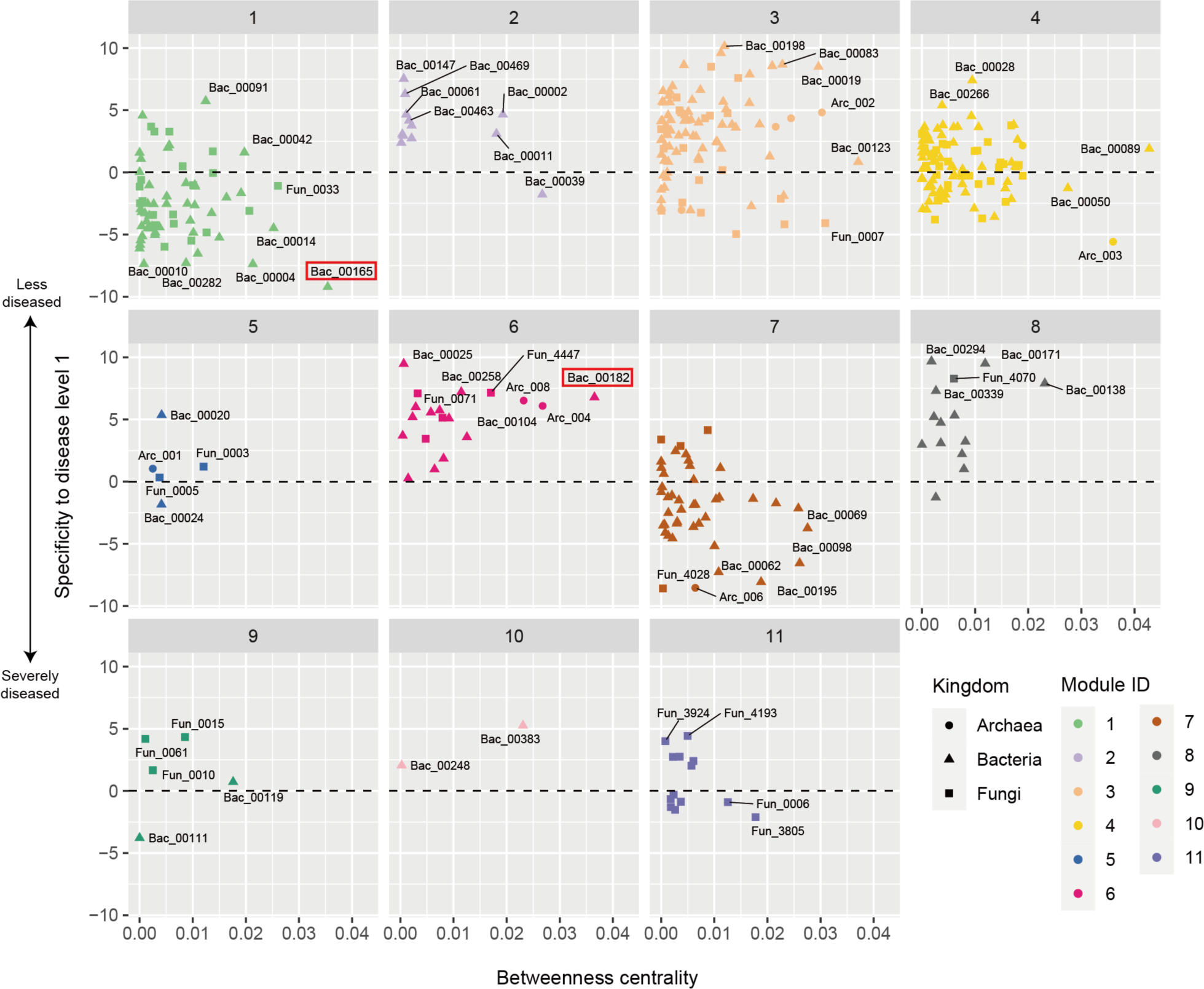
Properties of the microbe-to-microbe network modules. For each network module, specificity to the minimal crop disease level (disease level 1) is shown for each prokaryote/fungal OTU along the vertical axis. Betweenness centrality, which measures the extent to which an OTU is located within the shortest paths connecting pairs of other nodes in a network, is shown along the horizontal axis. The OTUs mentioned in the main text are highlighted with red squares.

We further ranked microbial OTUs in terms of their topological roles in interlinking multiple network modules. We then found that OTUs linked with many other OTUs within modules were not necessarily placed at the topological positions interconnecting different modules (Fig. 6). In Module 6, which showed high specificity to the minimal crop disease level (Fig. 4), a bacterium distantly allied to the genus *Thermanaerothrix* (Bac_00258) was designated as a “within-module hub”, while a *Plectosphaerella* fungus (Fun_4447) showed a high “among-module connectivity” score (Table 4). Likewise, in Module 1, which consisted of many OTUs with negative associations with crop plant health (Fig. 4), a bacterium allied to the genus *Gemmatimonas* (Bac_00258) had the highest numbers of within-module links, while a *Curvularia* fungus (Fun_0043) was inferred to be an among-module hub (Table 5). The list of microbial OTUs placed at the interface of modules (OTUs with high among-module connectivity scores) involved a *Nitrosotenuis* archaeum, *Arenimonas*, *Arthrobacter*, and *Streptomyces* bacteria, and *Mortierella*, *Curvularia*, and *Trichoderma* fungi (Table 5).

**Fig. 6.**
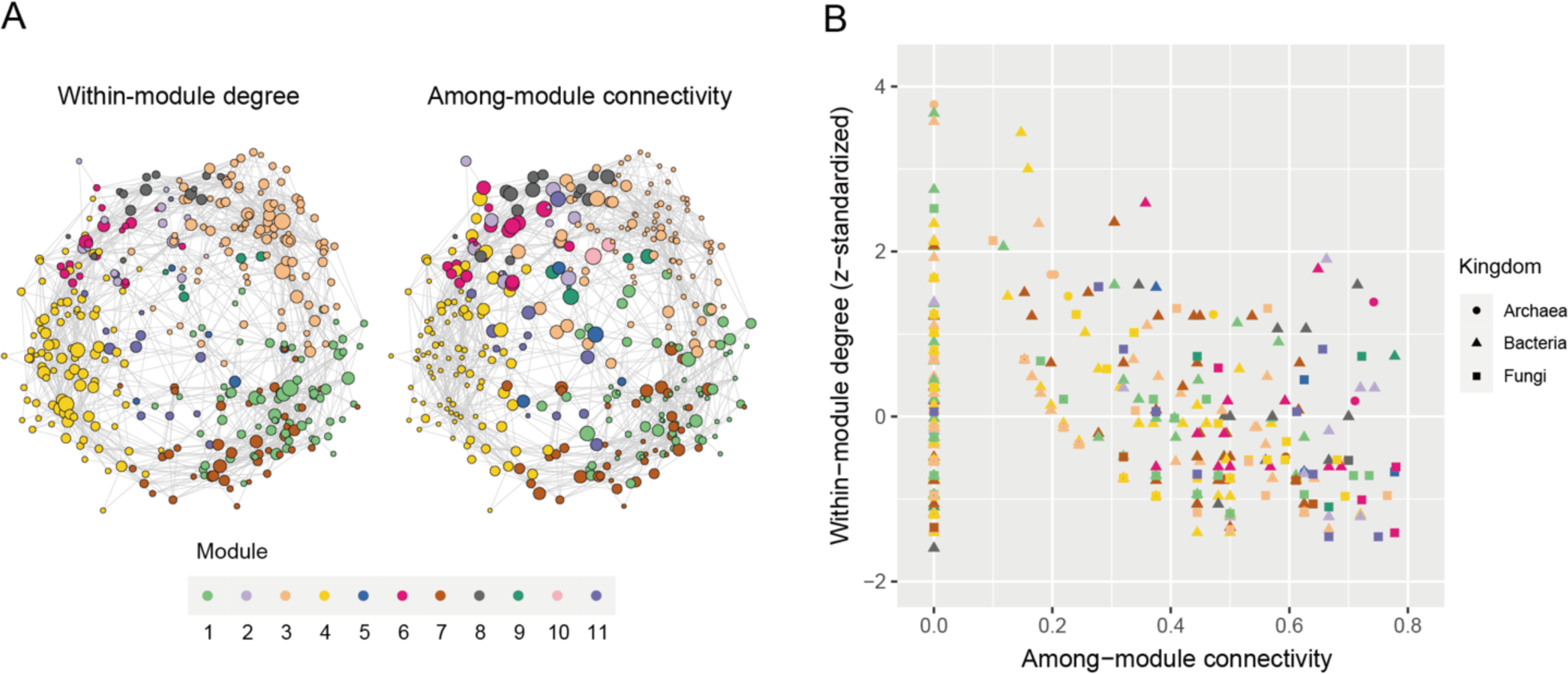
Topological roles of OTUs within and across network modules. (A) Position of potential hubs within the network. In each graph, node size roughly represents within-module degree (left) or among-module connectivity (right). (B) Network hub indices. For each OTU, within-module degree represents the number of the OTUs linked with the target OTU within a module (*z*-standardized). Among-module connectivity represents the extent to which an OTU interlinks OTUs belonging to different network modules. The prokaryotic/fungal OTU with the highest within-module degree or among-module connectivity in each of the modules 1, 2, 6, 7, and 8 (highlighted in the main text and Table 4) is indicated with its OTU ID. See Table 5 for the taxonomic profiles of the OTUs.

**Table 5.**
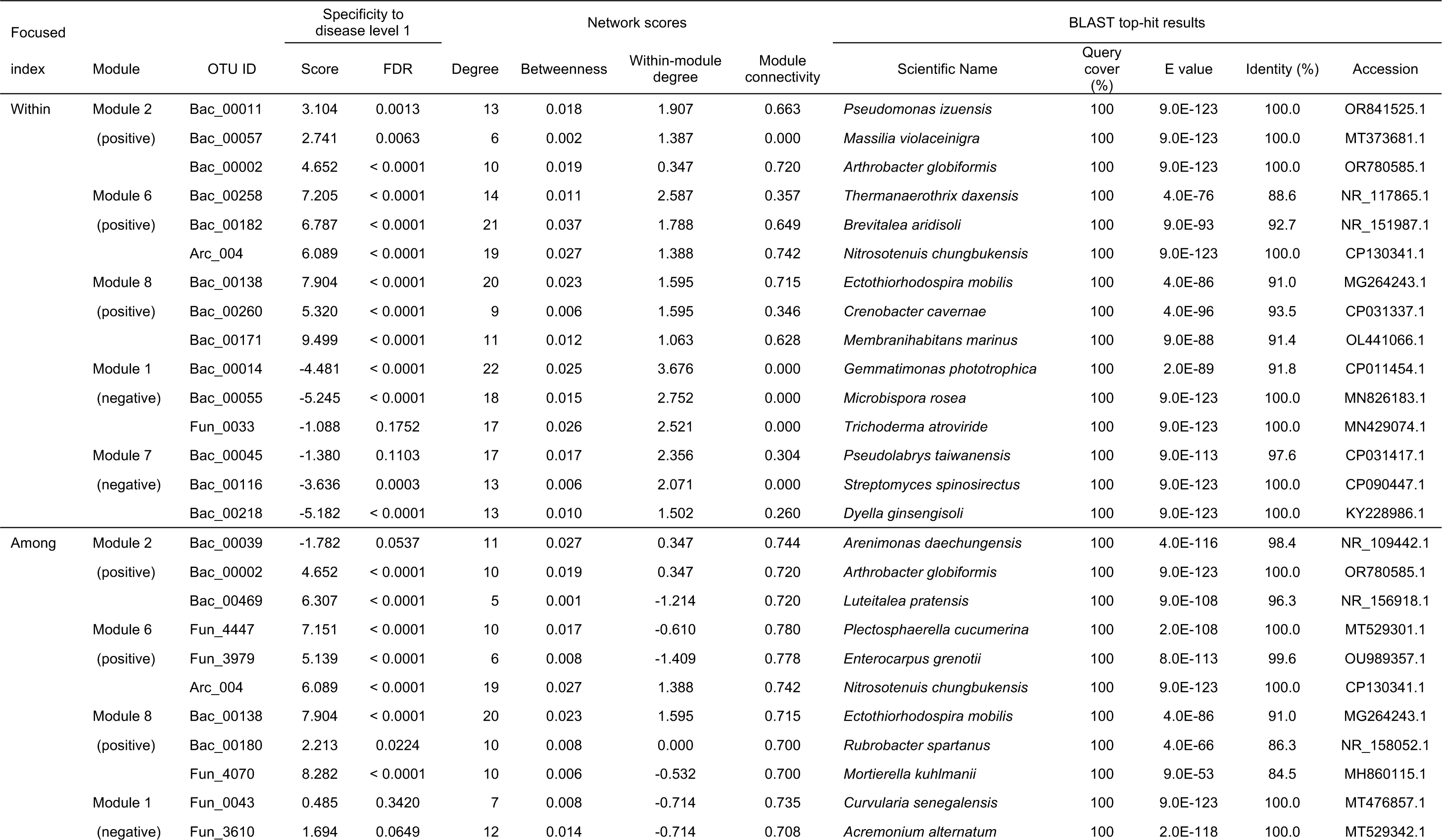
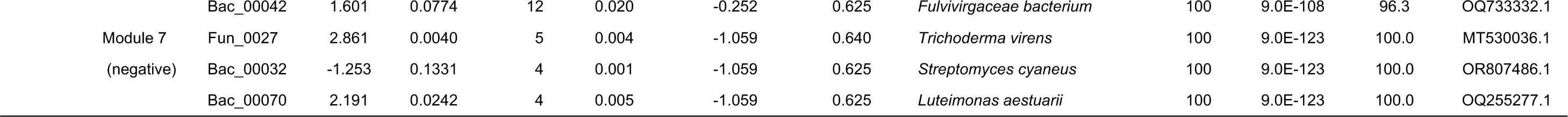
Within- and among-module hubs in the network. In each of the modules highlighted in Table 4 and Figure 6, the top-three OTUs with the highest within-module degree (*z*-standardized) or among-module connectivity are shown. See Data S3 for the full results.

## DISCUSSION

We here profiled the diversity of agroecosystem microbiome structure across a latitudinal gradient from cool-temperate to subtropical regions based on the analysis of > 2,000 soil samples. As partially reported in previous studies comparing microbiome compositions across broad geographic ranges (15, 22), prokaryotic and fungal community structure varied depending on season, crop plant species, former crop identity, and background soil categories (Fig. 2A; Table 1). In addition, soil chemical properties such as pH, electrical conductivity, and C/N ratio as well as the prokaryote/fungus abundance ratio significantly explained variation in microbiome structure (Table 1). In contrast, available phosphorus concentrations had significant effects on neither prokaryotic nor fungal communities in the multivariate model (Table 1), suggesting that nitrogen cycles rather than phosphorous ones are more tightly linked with microbiome structure. The integration of the microbiome datasets with agricultural field metadata allowed us to perform statistical tests of potential relationship between microbiome structure and agroecosystem performance (Fig. 2B; Table 2). A series of OTU-level analyses further highlighted taxonomically diverse prokaryotes and fungi showing strong positive or negative associations with crop health status (Figs. 3 and S4; Table 3).

We then examined how these microbes differing in associations with crop disease/health status form a network of coexistence. The architecture of the network involving diverse archaeal, bacterial, and fungal OTUs was highly structured, being partitioned into 11 modules (Fig. 4A). Intriguingly, the network modules varied considerably in constituent microbes’ associations with crop disease levels (Fig. 4B). This result suggests that sets of microbes can be used to design soil microbiomes with crop-disease-suppressive functions. Among the detected modules, Modules 2, 6, and 8 were of particular interest with regard to the assembly of microbial OTUs positively associated with crop health status (Figs. 4 and 5). In contrast, Modules 1 and 7 were constituted mainly by microbial OTUs negatively associated with plant health (Fig. 4B). In particular, Module 7 was characterized by the presence of a notorious plant pathogenic fungus, *Fusarium oxysporum* [(41, 42); but see (43) for diversity of their impacts on plants]. All these modules included both prokaryotes and fungi (Fig. S8; Data S3), illuminating the importance of inter-kingdom interactions (3, 33). The presence of microbial species sets differing in plant-associated ecological properties suggests that keeping specific sets of compatible prokaryotes and fungi is essential for maximizing the stability of agricultural production (3).

The analysis of network architecture further allowed us to explore core or hub species within the microbial network (Fig. 6). Because the microbes highlighted with the examined network indices occupy key positions interconnecting many other microbes (44), their increase/decrease is expected to have profound impacts on whole community processes (3, 32, 33). In particular, control or manipulation of microbes located at the central positions interlinking different network modules (40) (i.e., microbes with high among-module connectivity; Fig. 6B) may trigger drastic shifts in microbial community structure between disease-promotive and disease-suppressive states (3). The candidate list of such core species involved an ammonium-oxidizing archaeum (*Nitrosotenuis*) (45), an antibiotics-producing bacterium (*Streptomyces*) (46), a prevalent soil fungus (*Mortierella*) (47, 48), a potentially mycoparasitic fungus (*Trichoderma*) (49, 50), and fungi allied to plant pathogenic clades [*Curvularia* and *Plectosphaerella* (anamorph = *Fusarium*)] (51, 52) (Table 5). Given that many of the bacterial and fungal taxa listed above are culturable, experimental studies examining their ecological roles are awaited. Specifically, it would be intriguing to test whether substantial shifts in soil microbiome structure and functions can be caused by the introduction of those among-module hub microbes.

Although the dataset across a latitudinal gradient provided an opportunity for gaining bird’s-eye insights into the structure and potential functions of soil microbiomes, the results should be interpreted carefully with the recognition of potential methodological shortcomings and pitfalls. First, the approach of geographic comparison *per se* does not give a firm basis for deciphering microbial community dynamics. To gain fundamental insights into microbiome dynamics, we need to perform time-series monitoring (53–55) of soil prokaryotic and fungal community compositions. Second, information of microbial communities alone does not provide comprehensive insights into agroecosystem soil states. Given that soil ecosystem processes are driven not only by microbes but also by nematodes, arthropods, earthworms, and protists (56–59), simultaneous analyses of all prokaryotic and eukaryotic taxa (60, 61) will help us infer whole webs of biological processes. Third, meta-analyses of agroecosystem performance across diverse crop fields require utmost care because there is no firm criterion commonly applicable to different crop plant species or different pest/pathogen species. As implemented in this study, effects of such difference may be partially controlled by including them as random variables in GLMMs (Table 2). Nonetheless, local-scale analyses targeting specific crop plant species and disease symptoms (Fig. 3; Table 3; Data S2) are necessary to gain reliable inferences of potential microbial functions. Fourth, along with the potential pitfall discussed above, network modules can differ not only in properties related to crop disease/health status but also in those associated with crop plant identity or cropland management (Figs. S6-7). Again, findings in broad-geographic-scale analyses need to be supplemented by insights from local-scale observations (Fig. 3). Fifth, amplicon sequencing approaches provide only indirect inference of biological functions. With the current capacity of sequencing and bioinformatic technologies, it is hard to assemble tens of thousands of microbial genomes based on the analysis of thousands of environmental samples. Furthermore, due to the paucity of the information of fungal ecology and physiology, it remains difficult to annotate high proportions of genes within fungal genomic data. Nonetheless, with the accumulation of methodological breakthroughs, shotgun sequencing of soil microbiomes will deepen our understanding of agroecosystem processes (62–64). Sixth, in this study, full sets of metadata were not available for all the sequenced samples, inevitably decreasing the number of samples examined in some statistical modeling. Although substantial efforts had been made to profile cropland soils in the national projects in which the soil samples were collected, continuous efforts are required to gain further comprehensive insights into agroecosystem structure and functions.

Expanding the comparative microbiome analysis to different geographic regions and agroecosystem management practices will contribute to a more comprehensive understanding of microbiome structure and function. For example, comparison with soil agroecosystems in lower-latitudinal or higher-latitudinal regions or meta-analyses covering multiple continents will provide further comprehensive knowledge of the diversity of microbiome structure. In addition to extensions towards broader geographic ranges, those towards diverse agroecosystem management are of particular importance. Given that our samples were collected mainly from croplands managed with conventional agricultural practices, involvement of soil samples from regenerative or conservation agricultural fields (65–68) will reorganize our understanding of relationship between microbiome compositions and functions. In conclusion, this data-driven research lays the groundwork for understanding fundamental mechanisms in soil ecosystems, offering innovative strategies for the design of sustainable agriculture.

## MATERIALS AND METHODS

### Soil samples and metadata

Over research projects of National Agricultural and Food Research Organization (NARO), which were carried out through five national research programs funded by Ministry of Agriculture, Forestry and Fisheries, 2,903 rhizosphere/bulk soil samples were collected from conventional agricultural fields across the Japan Archipelago from January 23, 2006, to July 28, 2014 (Data S1). When the latitude and longitude of the sampling positions were round to one decimal place, 42 research sites were distinguished. Across the metadata of the 2,903 samples, the information of 19 crop plants, 34 former crop plants (including “no crop”), 13 soil taxonomic groups (e.g., “Andosol”), 60 experimental/research purposes (e.g., “soil comparison between organic and conventional management”) was described. Likewise, the metadata included the information of dry soil pH, electrical conductivity, carbon/nitrogen (C/N) ratio, and available phosphorous concentration from, 2,830, 2,610, 2,346, and 2,249 samples, respectively. In addition, the information of the severity of crop plant disease was available for 1,472 samples (tomato, 637 samples; Chinese cabbage, 336 samples; eggplant, 202 samples; celery, 97 samples; Broccoli, 96 samples, etc.). The values of the proportion of diseased plants or disease severity index (69) was normalized within the ranges from 0 to 100, and they were then categorized into five levels (level 1, 0-20; level 2, 20-40; level3; 40-60; level 4, 60-80; level 5, 80-100). The plant pathogens examined in the disease-level evaluation were *Colletotrichum gloeosporioides* on the strawberry, *Fusarium oxysporum* on the celery, the lettuce, the strawberry, and the tomato, *Phytophthora sojae* on the soybean, *Plasmodiophora brassicae* on Cruciferae plants, *Pyrenochaeta lycopersici* on the tomato, *Pythium myriotylum* on the ginger, *Ralstonia solanacearum* on the eggplant and the tomato, and *Verticillium* spp. on Chinese cabbage. For continuous variables within the metadata, emergent outliers (mean + 5 SD) were converted into “NA” in the data matrix used in the following statistical analyses as potential measurement/recording errors. Unrealistic electrical conductivity records (> 20) were converted into “NA” as well.

At each sampling position, five soil sub-samples collected from the upper layer (0-10 cm in depth) at five points (ca. 100 g each) were mixed. The mixed soil sample (ca. 500 g) was then sieved with 2-mm mesh in the field. The samples were stored at -20 °C until DNA extraction. In laboratory conditions, 0.4 g of soil (fresh weight) was subjected to DNA extraction with FastDNA SPIN Kit for Soil (Q-BioGene).

### DNA amplification and sequencing

Profiling of soil microbial biodiversity was performed by targeting archaea, bacteria, and fungi. For the amplification of the 16S rRNA V4 region of archaea and bacteria (prokaryotes), the set of the forward primer 515f (5’-GTG YCA GCM GCC GCG GTA A -3’) and the reverse primer 806rB (5’-GGA CTA CNV GGG TWT CTA AT -3’) were used as described elsewhere (54). The primers were fused with 3–6-mer Ns for improved Illumina sequencing quality and Illumina sequencing primers. PCR was performed using KOD ONE PCR Master Mix (TOYOBO, Osaka) with the temperature profile of 35 cycles at 98 °C for 10 seconds (denaturation), 55 °C for 5 seconds (annealing of primers), and 68 °C for 30 seconds (extension), and a final extension at 68 °C for 2 minutes. The ramp rate through the thermal cycles was set to 1 °C/sec to prevent the generation of chimeric sequences. In the PCR, we added five artificial DNA sequence variants with different concentrations (i.e., standard DNA gradients; 1.0 ×10^-4^, 5.0 ×10^-5^, 2.0 ×10^-5^, 1.0 ×10^-5^, and 5.0 ×10^-6^ nM; Table S1) to the PCR master mix solution as detailed elsewhere (54). By comparing the number of sequencing reads between the artificial standard DNA and real prokaryotic DNA, the concentration of prokaryotic 16S rRNA genes in template DNA samples were calibrated (54).

In addition to the prokaryotic 16S rRNA region, the internal transcribed spacer 1 (ITS1) region of fungi was amplified using the set of the forward primer ITS1F_KYO1 (5’-CTH GGT CAT TTA GAG GAA STA A -3’) and the reverse primer ITS2_KYO2 (5’ – TTY RCT RCG TTC TTC ATC - 3’) (70). PCR was performed using the Illumina-sequencing fusion primer design mentioned above with the temperature profile of 35 cycles at 98 °C for 10 seconds, 53 °C for 5 s seconds, and 68 °C for 5 seconds, and a final extension at 68 °C for 2 minutes (ramp rate = 1 °C/sec). Newly designed artificial sequence variants (1.0 ×10^-5^, 7.0 ×10^-6^, 5.0 ×10^-6^, 2.0 ×10^-6^, and 1.0 ×10^-6^ nM; Table S1) were added to the PCR master mix as standard DNA gradients for the calibration of the ITS sequence concentrations in the template DNA samples.

The PCR products of the prokaryotic 16S rRNA and fungal ITS1 regions were respectively subjected to the additional PCR step for linking Illumina sequencing adaptors and 8-mer sample identifier indexes with the amplicons. The temperature profile in the PCR was 8 cycles at 98 °C for 10 seconds, 55 °C for 5 seconds, and 68 °C for 5 seconds, and a final extension at 68 °C for 2 minutes. The PCR products were then pooled for each of the 16S rRNA and fungal ITS1 regions after a purification/equalization process with the AMPureXP Kit (Beckman Coulter, Inc., Brea). Primer dimers, which were shorter than 200 bp, were removed from the pooled library by supplemental purification with AMpureXP: the ratio of AMPureXP reagent to the pooled library was set to 0.8 (v/v) in this process. The sequencing libraries of the two regions were processed in an Illumina MiSeq sequencer (10% PhiX spike-in). Because the quality of forward sequences is generally higher than that of reverse sequences in Illumina sequencing, we optimized the MiSeq run setting in order to use only forward sequences. Specifically, the run length was set 271 forward (R1) and 31 reverse (R4) cycles to enhance forward sequencing data: the reverse sequences were used only for discriminating between prokaryotic 16S and fungal ITS1 sequences in the following bioinformatic pipeline.

### Bioinformatics

In total, 23,573,405 sequencing reads were obtained in the Illumina sequencing (16S rRNA, 11,647,166 sequencing reads; ITS, 11,926,239 sequencing reads). The raw sequencing data were converted into FASTQ files using the program bcl2fastq 1.8.4 distributed by Illumina. For each of the 16S rRNA and fungal ITS1 regions, the output FASTQ files were demultiplexed using Claident v0.9.2022.01.26 (71). The sequencing data were deposited to DNA Data Bank of Japan (DDBJ) (Bioproject accession no.: PSUB018361). The removal of low-quality sequences and OTU inferences were done using DADA2 (72) v1.17.5 of R v3.6.3. The mean number of filtered sequencing reads obtained per sample was 3,949 and 4,075 for the prokaryotic and fungal datasets, respectively. The amplicon sequence variants (ASVs) obtained from the DADA2 pipeline were clustered using the vsearch v2.21.1 program (73) with the 98% and 97% cutoff sequence similarity for prokaryotes and fungi, respectively. Taxonomic annotation of the obtained prokaryotic and fungal OTUs were conducted based on the SILVA 138 SSU (74) and the UNITE all_ 25.07.2023 (75) databases, respectively, with the assignTaxonomy function of DADA2. The OTUs that were not assigned to the domain Archaea/Bacteria and the kingdom Fungi were removed from the 16S rRNA and ITS1 datasets, respectively. For each target organismal group (prokaryotes and fungi), we then obtained a sample × OTU matrix, in which a cell entry depicted the number of sequencing reads of an OTU in a sample. The samples with less than 1,000 reads were discarded from the matrices. The number of reads was insufficient for comprehensively profiling rare microbial species, which are often targets of soil microbiome studies. However, because data matrices including numerous rare OTUs could not be subjected to the computationally intensive ecological analyses detailed below even if we used supercomputers, we focused on major components of soil prokaryotic and fungal biomes. In other words, our purpose here was to extract major components of agroecosystem soil microbiomes across the Japan Archipelago, thereby finding core microbiome properties associated with disease-suppressive and disease-susceptible agroecosystems. For the sample × OTU matrix, centered log-ratio (CLR) transformation (76–78) was performed using the ALDEx2 v1.35.0 package (79) of R.

In total, prokaryotic and fungal community data were obtained for 2,676 and 2,477 samples, respectively. For fungal OTUs, putative functional groups (e.g., “plant pathogen”) were inferred using the program FungalTraits (80). The estimation of DNA concentrations of the prokaryotic 16S rRNA and fungal ITS regions was performed, respectively, based on the calibration with the standard DNA gradients (artificial DNA variants introduced to the PCR master mix solutions) using the bioinformatic pipeline detailed elsewhere (54).

### Calculation of prokaryote/fungus ratio

Based on the estimated concentrations of prokaryotic 16S rRNA and fungal ITS sequences in template DNA solutions, we calculated the ratio of prokaryotic DNA concentrations to fungal DNA concentrations in respective samples (prokaryote/fungus ratio) as follows:

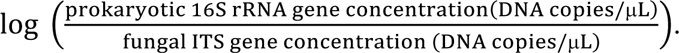

Although potential variation in DNA extraction skills of researchers might affect absolute DNA concentrations in the template DNA solutions, balance between prokaryotic and fungal DNA in each template DNA sample could be used as a reliable measure. The DNA-metabarcoding-based approach of estimating prokaryote/fungus ratio has methodological advantage over quantitative-PCR-based approaches. Specifically, the former approach allows us to eliminate effects of nonspecific PCR amplification based on DNA sequencing data, while the latter is affected by “contamination” of nontarget amplicons (e.g., plastid DNA in 16S rRNA sequencing and plant DNA in ITS sequencing).

### Microbiome structure and crop disease prevalence

For each of the prokaryotic and fungal datasets, permutational analyses of variance (PERMANOVA) (81) were performed to examine associations between family-level community compositions and variables in the metadata. Two types of PERMANOVA models were constructed based on the Euclid distance (*β*-diversity) calculated for the CLR-transformed datasets (1,000 iterations). Specifically, one is constituted by categorical explanatory variables (crop plant, former crop plant, soil taxonomy, research site, and sampling month), while the other included continuous explanatory variables (soil pH, electrical conductivity, C/N ratio, and available phosphorous concentration, prokaryote/fungus ratio, latitude, and longitude).

To reduce the dimensions of the community compositional data, a principal component analysis (PCA) was performed based on the Euclid distance data mentioned above. For each PCA axis (axes 1 to 5) in each of the prokaryotic and fungal analyses, Pearson’s correlation with each chemical environmental factor (soil pH, electrical conductivity, C/N ratio, and available phosphorous concentration) was calculated.

We then evaluated how community structure of prokaryotes and fungi were associated with crop disease. For each of the prokaryotic and fungal datasets, a generalized linear mixed model (GLMM) of crop-disease level (disease level 1 vs. disease levels 2-5) was constructed by including the PCoA axes 1 to 5 as fixed effects. Sampling month and the identity of crop plant species and experimental/research purposes in the metadata were set as random effects. A logit-link function and binomial errors was assumed in the GLMM after converting the response variable into a binary format [disease level 1 (= 1) vs. disease levels 2–5 (= 0)]. The analysis was performed with the “glmer” function of the R lme4 package (82).

### Microbes associated with crop disease/health status

For each microbial OTU constituting the modules, we evaluated specificity of occurrences in samples differing in crop disease levels based on a randomization analysis. For the calculation, the original sample × OTU matrices of prokaryotes and fungi were respectively rarefied to 1,000 reads per sample, being merged into an input data matrix. Within the combined sample × OTU matrix, samples were categorized into the two crop disease levels (disease level 1 vs. disease levels 2-5). Mean read counts across samples displaying each of the two disease levels were then calculated for each OTU. Meanwhile, mean read counts for respective disease levels were calculated as well for randomized matrices, in which disease labels of the samples were shuffled (10,000 permutations). For *i*-th OTU, standardized specificity to disease level *j* (*s_ij_*) was obtained as follows:

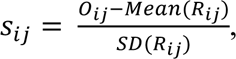

where *O_ij_* and *R_ij_* is the mean read counts of *i*-th OTU across disease-level-*j* samples in the observed and randomized matrices, respectively, and *Mean*(*R_ij_*) and *SD*(*R_ij_*) indicate mean and standard deviation across the randomized matrices. The *P* values obtained based on the randomization analysis were adjusted with the Benjamini-Hochberg method [i.e., false discovery rate (FDR)]. The relationship between the standardized specificity index and FDR is shown in Figure S4. This randomization approach was also applied to the analyses of each OTU’s specificity to crop plant identity and that to experimental/research purpose identity (Figs. S6-7).

The specificity of microbial OTUs to crop disease levels was performed as well at the local scale. Specifically, in each of the six sub-datasets representing unique combinations of research sites, crop plant species, and experimental/research purposes, the abovementioned randomization analysis was performed: each sub-dataset included 69 to 198 soil samples (Data S2). For the OTUs showing exceptionally strong specificity to the minimal crop disease level (standardized specificity score ≥ 6.0; FDR < 0.0001), supplemental analyses of generalized linear models (GLMs) were conducted. In each GLM of crop disease/health status (disease level 1 vs. disease levels 2-5) with a logit-link function with binomial errors, the relative abundance of a target OTU was included as an explanatory variable.

### Microbe-to-microbe network

To infer potential interactions between microbial OTUs, the algorithm of sparse inverse covariance estimation for ecological association inference (SPIEC-EASI) was applied based on the Meinshausen-Bühlmann (MB) method as implemented in the SpiecEasi package (39) of R. In total, 2,305 soil samples from which both prokaryotic and fungal community data were available were subjected to the analysis. Note that CLR-transformation was performed internally with the “spiec.easi” function. The network inference based on co-occurrence patterns allowed us to detect pairs of microbial OTUs that potentially interact with each other in facilitative ways and/or those that might share ecological niches (e.g., preference for edaphic factors). Because estimation of co-occurrence patterns was not feasible for rare nodes, the prokaryotic and fungal OTUs that appeared in more than 10 % of the sequenced samples were included in the input matrix of the network analysis. Network modules, within which closely associated OTUs were interlinked with each other, were identified with the algorithm based on edge betweenness (83) using the igraph package (84) of R. For each module in the inferred co-occurrence network, mean standardized specificity to disease level 1 were calculated across constituent OTUs.

To explore potential keystone microbes within the network, we scored respective OTUs on the basis of their topological positions. Among the indices used for evaluating OTUs, betweenness centrality (85), which measures the extent to which a given nodes (OTU) is located within the shortest paths connecting pairs of other nodes in a network, is commonly used to find hubs mediating flow of effects in a network. The network centrality scores were normalized as implemented in the igraph packages of R. In addition, by focusing on the above-mentioned network modules, we ranked OTUs based on their within-module degree and among-module connectivity (86). The former index is obtained as the number of nodes linked with a target node within a target network module, suggesting the topological importance of a node within the module it belongs to. The latter index represents the extent to which a node is linked with other nodes belonging to different network modules. Within-module degree was *z*-standardized (i.e., zero-mean and unit-variance) within each module, while among-module connectivity was defined to vary between 0 to 1. In addition to those indices for evaluating topological roles within a network, eigenvector centrality (87) was calculated for respective nodes.

### Data availability

The 16S rRNA and ITS sequencing data are available from the DNA Data Bank of Japan (DDBJ accession: DRA015491 and DRA015506). The microbial community data are deposited at our GitHub repository (https://github.com/hiro-toju/Soil_Microbiome_NARO3000).

### Code availability

All the R scripts used to analyze the data are available at the GitHub repository (https://github.com/hiro-toju/Soil_Microbiome_NARO3000).

## ACKNOWLEDGEMENTS

We are grateful to the reviewers whose comments improved the manuscript. We thank the SuperComputer System, Institute for Chemical Research, Kyoto University for the use of super computers. This work was financially supported by JST PRESTO (JPMJPR16Q6), JST FOREST (JPMJFR2048), JST PRESTO (JPMJCR23N5), Human Frontier Science Program (RGP0029/2019), JSPS Grant-in-Aid for Scientific Research (20K20586) and NEDO Moonshot Research and Development Program (JPNP18016) to H.T., JSPS Grant-in-Aid for Scientific Research (20K06820 and 20H03010) to K.S., and JSPS Fellowship to H.F.

