## Supplementary Figures for "Soil Prokaryotic and Fungal Biome Structures Associated with Crop Disease Status across the Japan Archipelago"

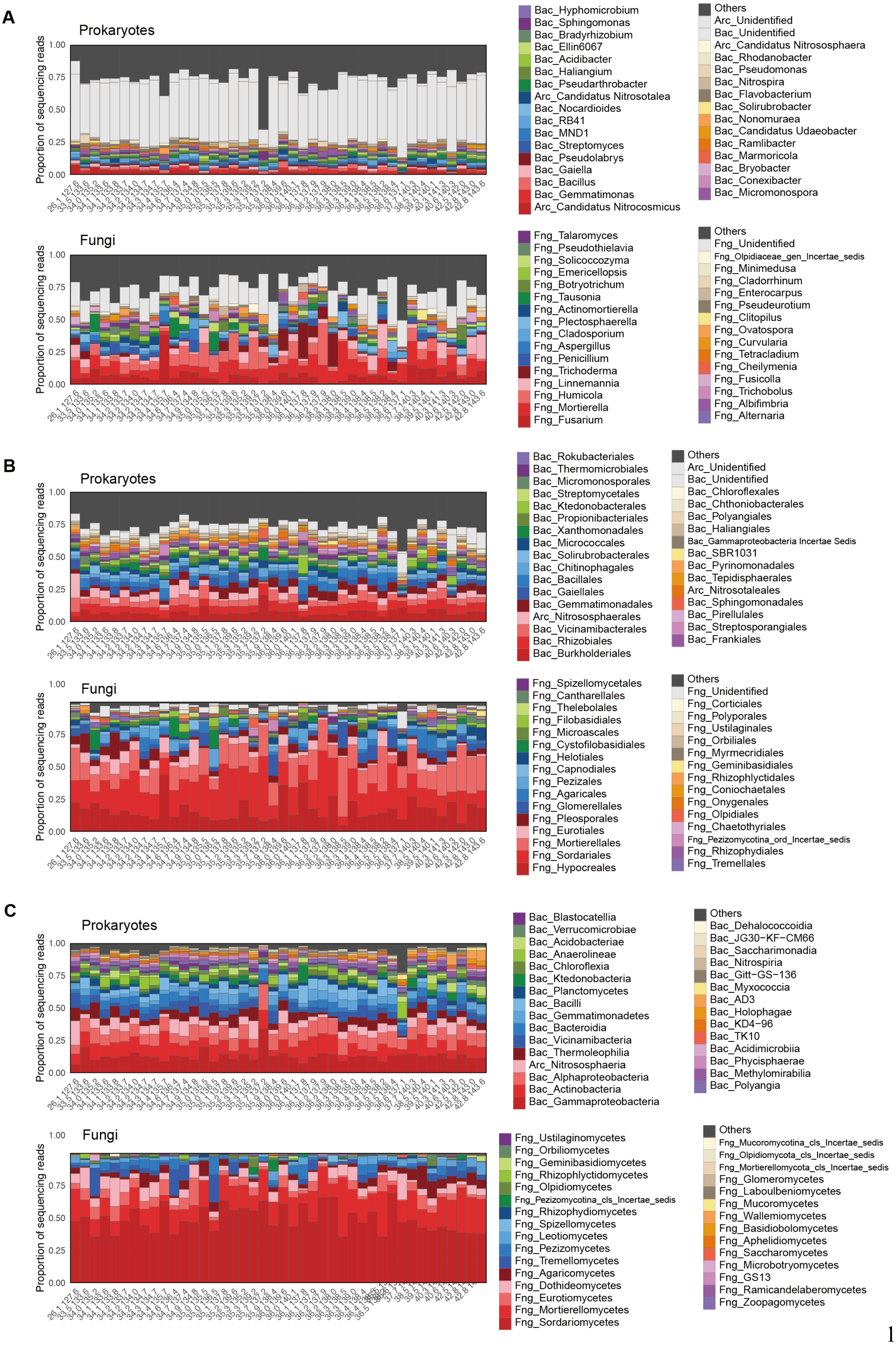
A
Prokaryotes
Proportion of sequencing reads
1.00
0.75
0.50
0.25
0.00
Bac_Hyphomicrobium
Bac_Sphingomonas
Bac_Bradyrhizobium
Bac_Ellin6067
Bac_Acidibacter
Bac_Haliangium
Bac_Pseudarthrobacter
Arc_Candidatus Nitrosotalea
Bac_Nocardioides
Bac_RB41
Bac_MND1
Bac_Streptomyces
Bac_Pseudolabrys
Bac_Gaiella
Bac_Bacillus
Bac_Gemmatimonas
Arc_Candidatus Nitrocosmicus
Others
Arc_Unidentified
Bac_Unidentified
Arc_Candidatus Nitrososphaera
Bac_Rhodanobacter
Bac_Pseudomonas
Bac_Nitrospira
Bac_Flavobacterium
Bac_Solirubrobacter
Bac_Nonomuraea
Bac_Candidatus Udaeobacter
Bac_Ramlibacter
Bac_Marmoricola
Bac_Bryobacter
Bac_Conexibacter
Bac_Micromonospora
Fungi
Proportion of sequencing reads
Fng_Talaromyces
Fng_Pseudothielavia
Fng_Solicoccozyma
Fng_Emericellopsis
Fng_Botryotrichum
Fng_Tausonia
Fng_Actinomortierella
Fng_Plectosphaerella
Fng_Cladosporium
Fng_Aspergillus
Fng_Penicillium
Fng_Trichoderma
Fng_Linnemannia
Fng_Humicola
Fng_Mortierella
Fng_Fusarium
Others
Fng_Unidentified
Fng_Olpidiaceae_gen_Incertae_sedis
Fng_Minimedusa
Fng_Cladorrhinum
Fng_Enterocarpus
Fng_Pseudeurotium
Fng_Clitopilus
Fng_Ovatospora
Fng_Curvularia
Fng_Tetracladium
Fng_Cheilymenia
Fng_Fusicolla
Fng_Trichobolus
Fng_Albifimbria
Fng_Alternaria
B
Prokaryotes
Proportion of sequencing reads
Bac_Rokubacteriales
Bac_Thermomicrobiales
Bac_Micromonosporales
Bac_Streptomycetales
Bac_Ktedonobacterales
Bac_Propionibacteriales
Bac_Xanthomonadales
Bac_Micrococcales
Bac_Solirubrobacterales
Bac_Chitinophagales
Bac_Bacillales
Bac_Gaiellales
Bac_Gemmatimonadales
Arc_Nitrososphaerales
Bac_Vicinamibacterales
Bac_Rhizobiales
Bac_Burkholderiales
Others
Arc_Unidentified
Bac_Unidentified
Bac_Chloroflexales
Bac_Chthoniobacterales
Bac_Polyangiales
Bac_Haliangiales
Bac_Gammaproteobacteria Incertae Sedis
Bac_SBR1031
Bac_Pyrinomonadales
Bac_Tepidisphaerales
Arc_Nitrosotaleales
Bac_Sphingomonadales
Bac_Pirellulales
Bac_Streptosporangiales
Bac_Frankiales
Fungi
Proportion of sequencing reads
Fng_Spizellomycetales
Fng_Cantharellales
Fng_Thelebolales
Fng_Filobasidiales
Fng_Microascales
Fng_Cystofilobasidiales
Fng_Helotiales
Fng_Capnodiales
Fng_Pezizales
Fng_Agaricales
Fng_Glomerellales
Fng_Pleosporales
Fng_Eurotiales
Fng_Mortierellales
Fng_Sordariales
Fng_Hypocreales
Others
Fng_Unidentified
Fng_Corticiales
Fng_Polyporales
Fng_Ustilaginales
Fng_Orbiliales
Fng_Myrmecridiales
Fng_Geminibasidiales
Fng_Rhizophlyctidales
Fng_Coniochaetales
Fng_Onygenales
Fng_Olpidiales
Fng_Chaetothyriales
Fng_Pezizomycotina_ord_Incertae_sedis
Fng_Rhizophydiales
Fng_Tremellales
C
Prokaryotes
Proportion of sequencing reads
Bac_Blastocatellia
Bac_Verrucomicrobiae
Bac_Acidobacteriae
Bac_Anaerolineae
Bac_Chloroflexia
Bac_Ktedonobacteria
Bac_Planctomycetes
Bac_Bacilli
Bac_Gemmatimonadetes
Bac_Bacteroidia
Bac_Vicinamibacteria
Bac_Thermoleophilia
Arc_Nitrososphaeria
Bac_Alphaproteobacteria
Bac_Actinobacteria
Bac_Gammaproteobacteria
Others
Bac_Dehalococcoidia
Bac_JG30−KF−CM66
Bac_Saccharimonadia
Bac_Nitrospiria
Bac_Gitt−GS−136
Bac_Myxococcia
Bac_AD3
Bac_Holophagae
Bac_KD4−96
Bac_TK10
Bac_Acidimicrobiia
Bac_Phycisphaerae
Bac_Methylomirabilia
Bac_Polyangia
Fungi
Proportion of sequencing reads
Fng_Ustilaginomycetes
Fng_Orbiliomycetes
Fng_Geminibasidiomycetes
Fng_Rhizophlyctidomycetes
Fng_Olpidiomycetes
Fng_Pezizomycotina_cls_Incertae_sedis
Fng_Rhizophydiomycetes
Fng_Spizellomycetes
Fng_Leotiomycetes
Fng_Pezizomycetes
Fng_Tremellomycetes
Fng_Agaricomycetes
Fng_Dothideomycetes
Fng_Eurotiomycetes
Fng_Mortierellomycetes
Fng_Sordariomycetes
Others
Fng_Mucoromycotina_cls_Incertae_sedis
Fng_Olpidiomycota_cls_Incertae_sedis
Fng_Mortierellomycota_cls_Incertae_sedis
Fng_Glomeromycetes
Fng_Laboulbeniomycetes
Fng_Mucoromycetes
Fng_Wallemiomycetes
Fng_Basidiobolomycetes
Fng_Aphelidiomycetes
Fng_Saccharomycetes
Fng_Microbotryomycetes
Fng_GS13
Fng_Ramicandelaberomycetes
Fng_Zoopagomycetes



1

**Fig. S1 |** Microbial taxonomic compositions at the genus, order, and class levels. (A) Genus-level taxonomic compositions of prokaryotes and fungi. See Figure 1A for the map of the research sites. (B) Order-level taxonomic compositions. (C) Class-level taxonomic compositions.

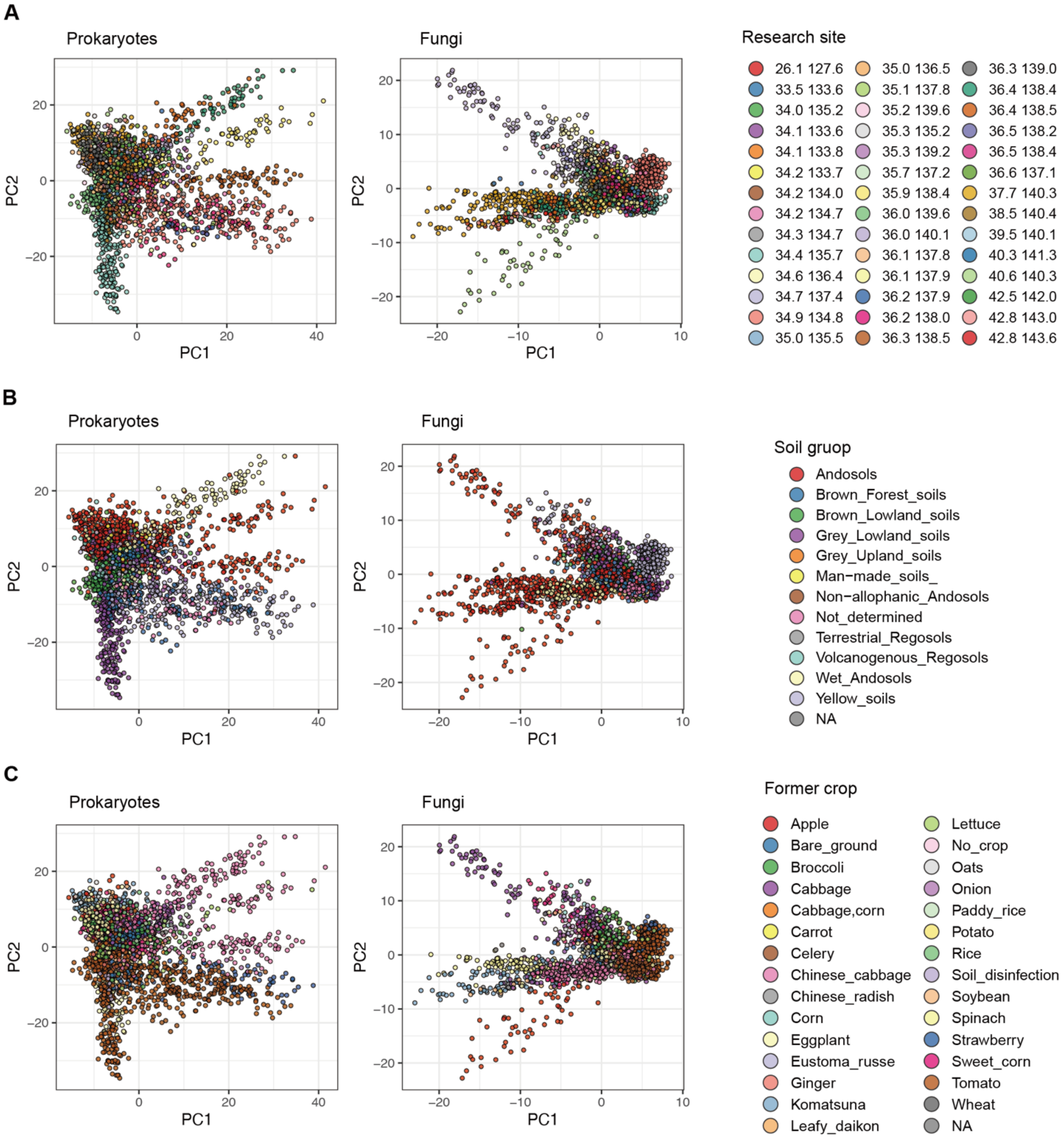


**Fig. S2 |** Community structure and metadata properties of the samples. (A) Prokaryote/fungal community structure and research-site profiles. Research sites are indicated by colors on the PCA surface of prokaryotic/fungal community structure. (B) Soil taxonomy profile. (C) Former crop plant.

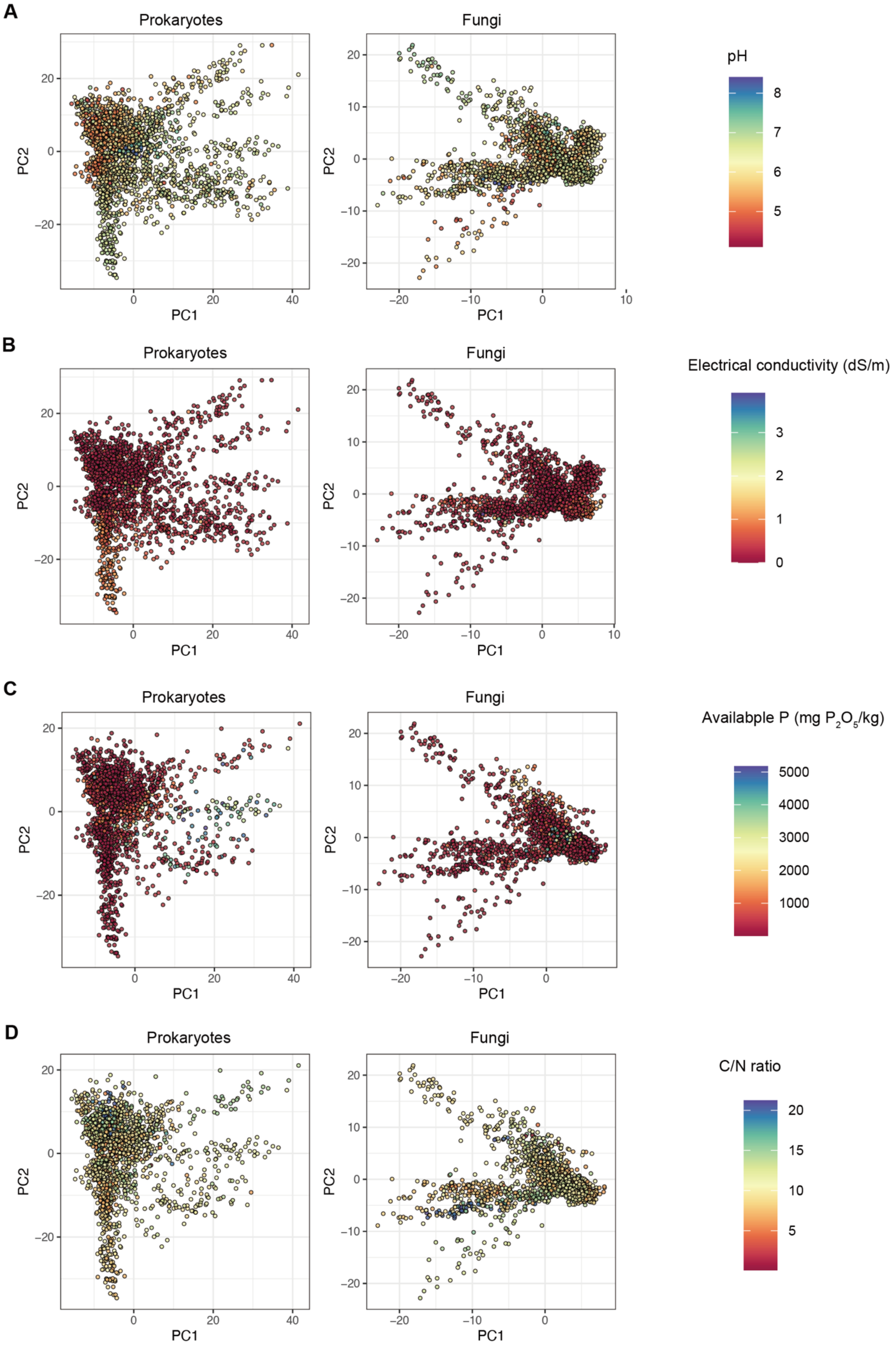
A
Prokaryotes
Fungi
pH
PC1
PC2
B
Electrical conductivity (dS/m)
C
Availabple P (mg P$_2$O$_5$/kg)
D
C/N ratio

13

**Fig. S3 |** Community structure and soil chemical properties. (A) Prokaryote/fungal community structure and soil pH of the samples. (B) Electrical conductivity. (C) Available phosphorous concentration. (D) Carbon to nitrogen ratio.

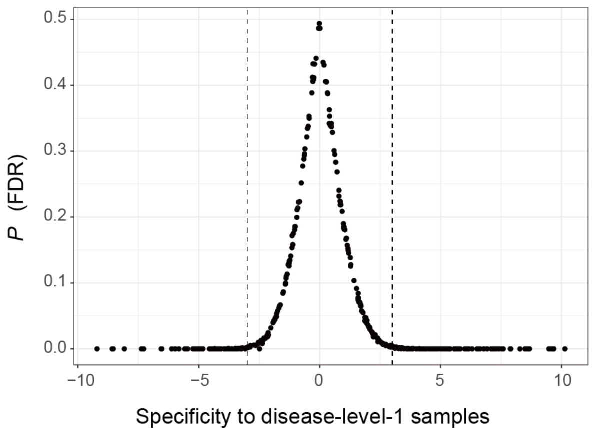


**Fig. S4 |** Index of specificity to crop disease levels. Based on a randomization approach, each OTU's specificity to samples differing in crop disease levels was evaluated. A higher value of the specificity index indicates that a microbial OTU displayed higher abundance in samples of the minimal crop disease level (disease level 1) than that expected by chance. Relationship between the standardized specificity index and false discovery rate (FDR) is shown.

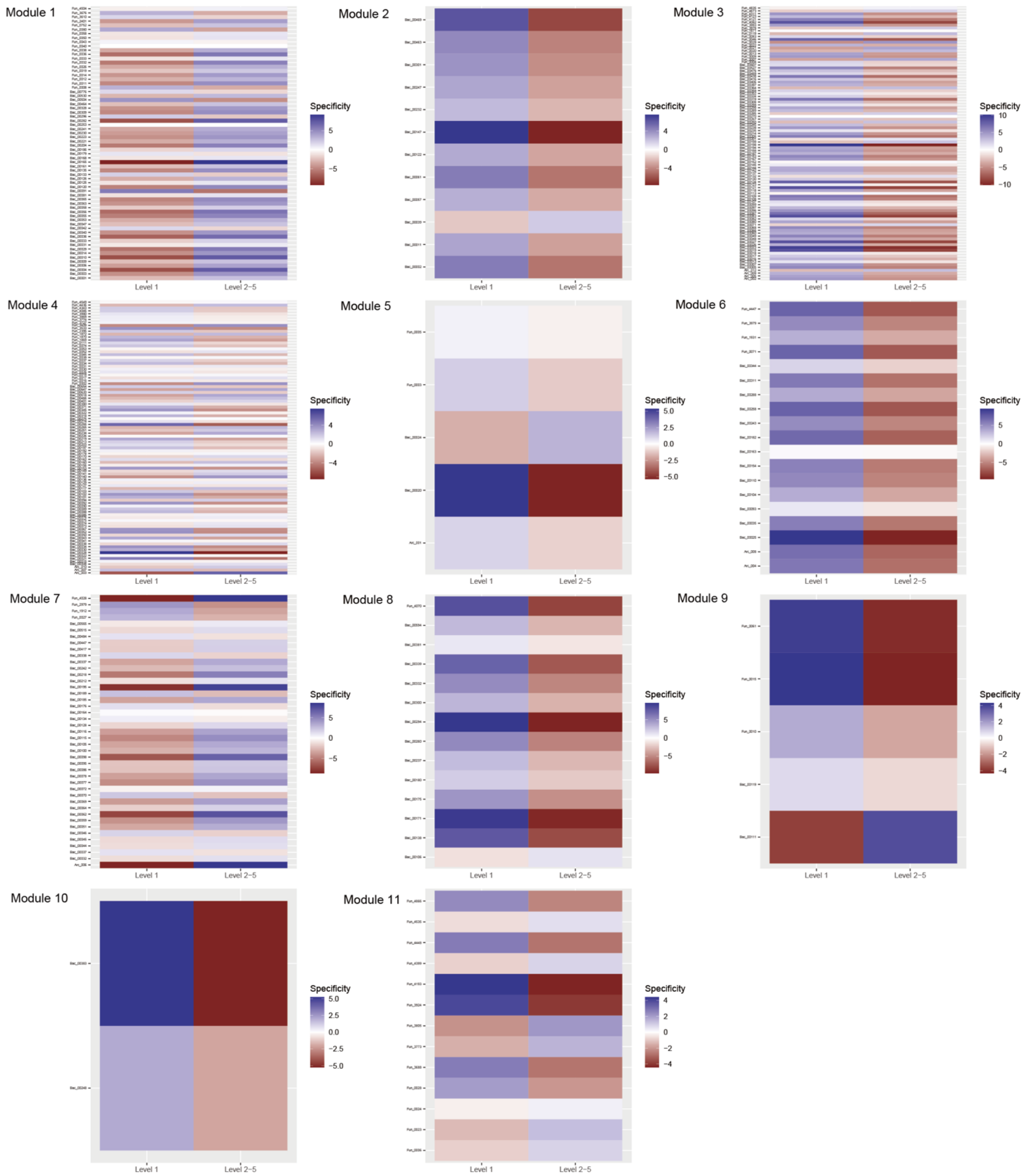


**Fig. S5 |** Each OTU's specificity to crop disease levels. Based on a randomization approach, each OTU's specificity to samples differing in crop disease levels was evaluated. OTUs belonging to respective network modules (Fig. 4) are separately shown.

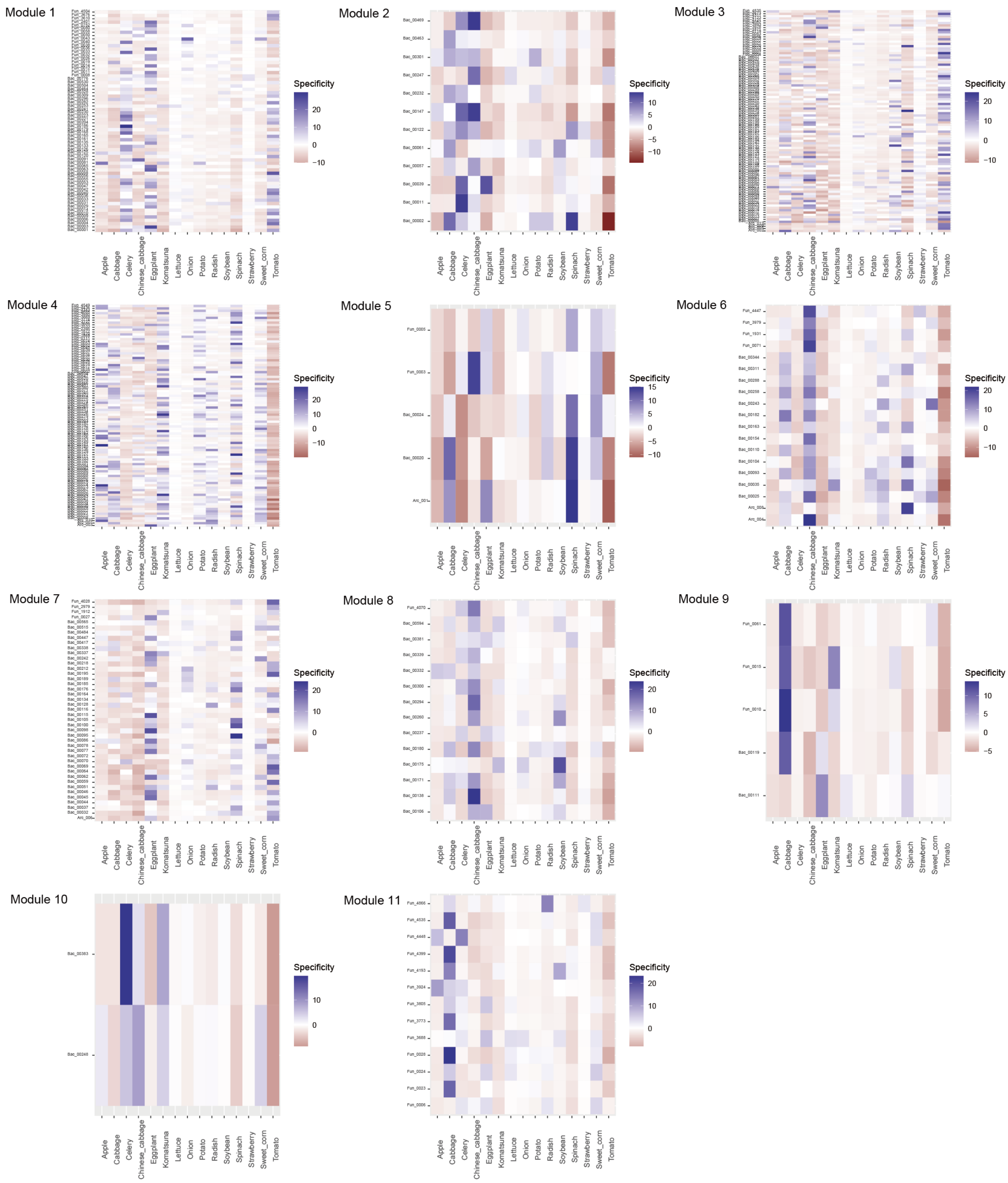


**Fig. S6 |** Each OTU's specificity to crop plant species. Based on a randomization approach, each OTU's specificity to samples differing in crop plant identity was evaluated. OTUs belonging to respective network modules (Fig. 4) are separately shown.

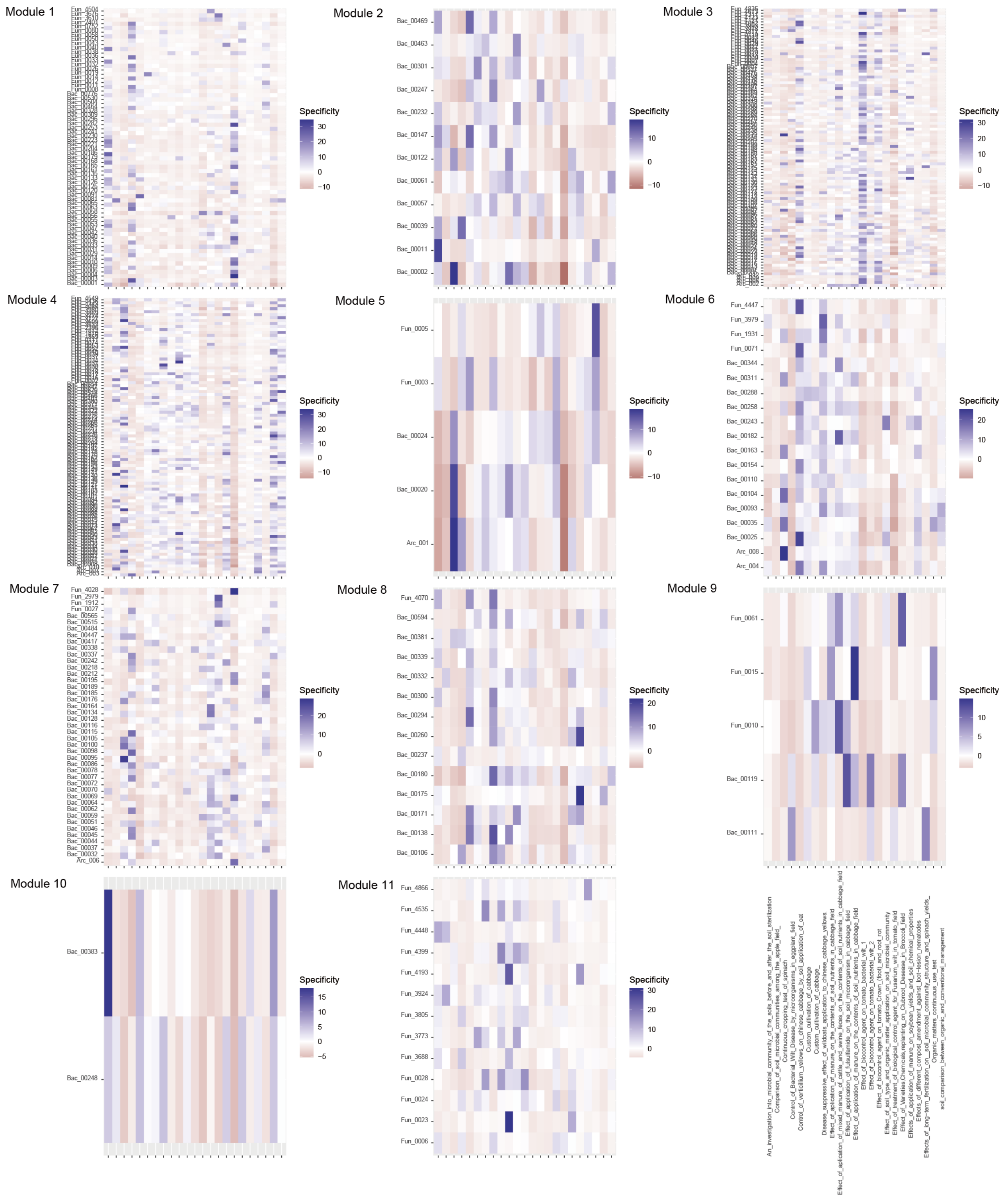


**Fig. S7 |** Each OTU's specificity to experimental/research purpose identity. (A) Standardized specificity to disease level ($s_{ij}$). Based on a randomization approach, each OTU's specificity to samples differing in experimental/research purpose identity was evaluated. OTUs belonging to respective network modules (Fig. 4) are separately shown.

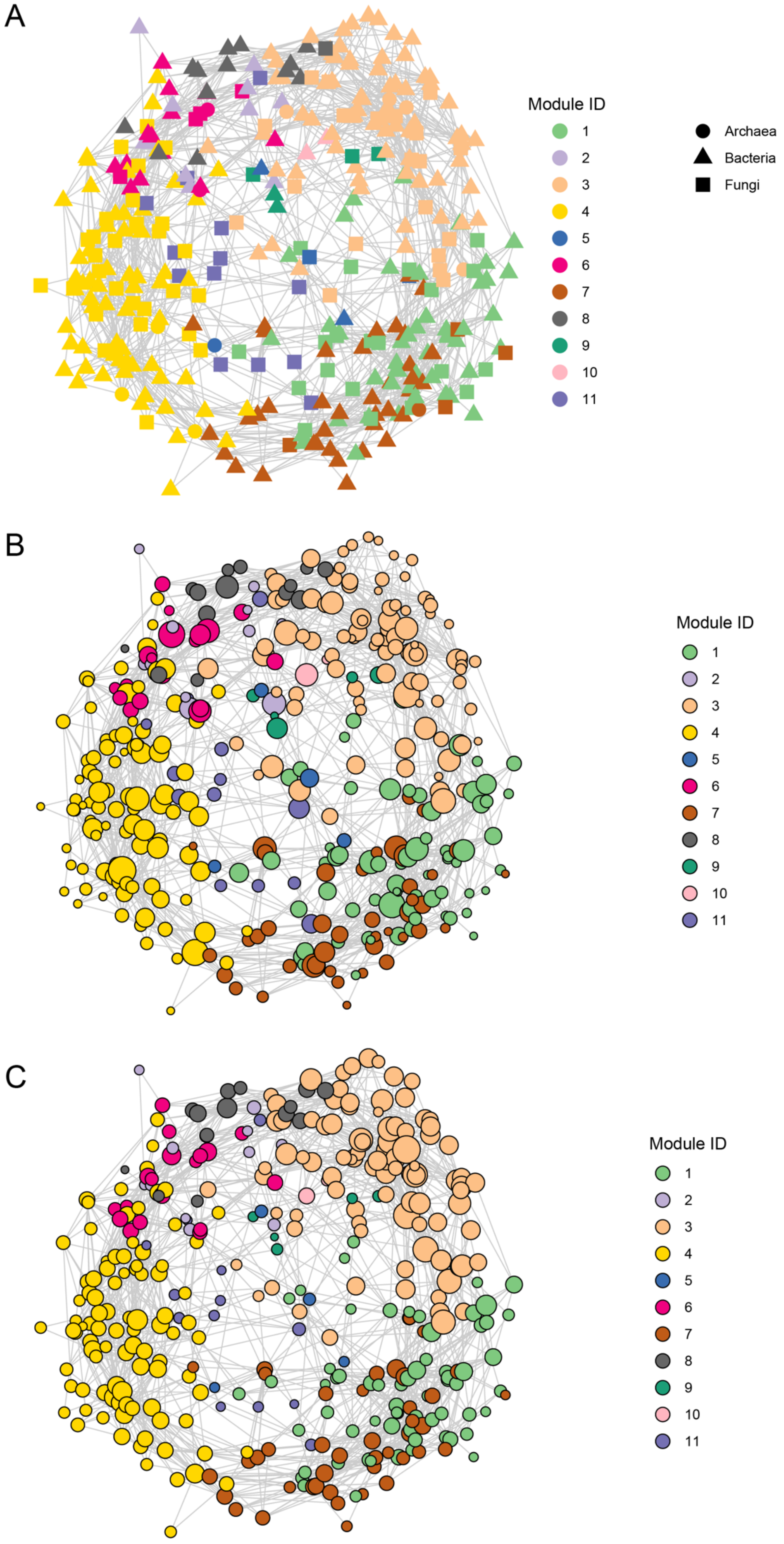
A
Module ID
1
2
3
4
5
6
7
8
9
10
11
Archaea
Bacteria
Fungi
B
Module ID
1
2
3
4
5
6
7
8
9
10
11
C
Module ID
1
2
3
4
5
6
7
8
9
10
11

**Fig. S8 |** Network centrality scores. (A) Distribution of archaea, bacteria, and fungi within the co-occurrence network. (B) Each OTU's betweenness centrality score within the network. The node size roughly represents relative betweenness centrality scores. (C) Each OTU's eigenvector centrality score within the network. The node size roughly represents relative eigenvector centrality scores.
