## Supplementary Table for "Soil Prokaryotic and Fungal Biome Structures Associated with Crop Disease Status across the Japan Archipelago"

Table S1 | Artificial sequences used for the calibration of prokaryote/fungal DNA concentrations. The artificial sequences with defined concentrations were mixed with the PCR master mix solutions as standard DNA gradients. The artificial sequences consisted of primer annealing regions, conservative sequence regions adjacent to the primer regions, and random nucleotide sequences at the highly-variable region at the intermediate positions. The standard (artificial) sequences were distinguished in the bioinformatic pipeline and they were used for the estimation (calibration) of prokaryote 16S rRNA or fungal ITS DNA concentrations in template DNA samples as detailed elsewhere^1–3^.

Standard DNA variants used in the prokaryote analysis^2^.

>STD_pro1

GTGCCAGCAGCCGCGGTAAGACGGAGGGGGCTAGCGTTGTTCGGAATTACTGGGCGTAAAGAGAAGGTAGGCGGAAGCTGAAGTCATGTGTGAAAACGCCTGGCTTAACTTAGCTCAGGGTCGCTAAACTGGTTGGCTTGAGTGTGAACGAGGTCCTCGGAATTTTCTGTGTAGCGGTGAAATGCGTAGATATTAAGGCGAACACCTGCGGCGAAGGCACGGAGCTGGGGCAGGGCTGACGCTGAGGCGTTAAAGCGTGGGGAGCAAACAGGATTAGATACCCTGGTAGTCC

>STD_pro2

GTGCCAGCAGCCGCGGTAAGACGGAGGGGGCTAGCGTTGTTCGGAATTACTGGGCGTAAAGAGTATGTAGGCGGTAAAGGAAGTTACGAGTGAAATTACAGGGCTTAACCGATAAGTCGTGGCCAAAACTGGGAGCCTTGAGTAATCGAGAGGTGGGCGGAATTGGGTGTGTAGCGGTGAAATGCGTAGATATTCAAAGGAACACCGATCGCGAAGGCGGCCTCCTGGTTAGGTCCTGACGCTGAGGAACGAAAGCGTGGGGAGCAAACAGGATTAGATACCCTGGTAGTCC

>STD_pro3

GTGCCAGCAGCCGCGGTAAGACGGAGGGGGCTAGCGTTGTTCGGAATTACTGGGCGTAAAGAGTTAGTAGGCGGGCAATTAAGTTATAGGTGAAAAGTAATGGCTTAACTTCGCGAACTCCGGACAAACTGAGGTGCTTGAGCGTTAAAGAGGCTCCCGGAATTGAAGGTGTAGCGGTGAAATGCGTAGATATGTTTCGGAACACCTAATGCGAAGGCTCAAGTCTGGGTAACTGGTGACGCTGAGGTCACAAAGCGTGGGGAGCAAACAGGATTAGATACCCTGGTAGTCC

>STD_pro4

GTGCCAGCAGCCGCGGTAAGACGGAGGGGGCTAGCGTTGTTCGGAATTACTGGGCGTAAAGAGAGTGTAGGCGGCCACGTAAGTTCCCGGTGAAATCGAGCGGCTTAACGTGCTCCTCGCCCGAGAAACTGAAGCCCTTGAGCTCAGCCGAGGAACCCGGAATTACTTGTGTAGCGGTGAAATGCGTAGATATGTGTATGAACACCTCCAGCGAAGGCCGCACACTGGCACTCCACTGACGCTGAGGTTTAAAAGCGTGGGGAGCAAACAGGATTAGATACCCTGGTAGTCC

>STD_pro5

GTGCCAGCAGCCGCGGTAAGACGGAGGGGGCTAGCGTTGTTCGGAATTACTGGGCGTAAAGAGCCGGTAGGCGGCTCCCGAAGTCGGTAGTGAAATTCTGAGGCTTAACTAAAACCACGACATTGAAACTGGGTGTCTTGAGTTGATACGAGGCCAGTGGAATTGTGCGTGTAGCGGTGAAATGCGTAGATATAATAAGGAACACCTCCCGCGAAGGCTTTGGCCTGGCATGCCAGTGACGCTGAGGTGTTAAAGCGTGGGGAGCAAACAGGATTAGATACCCTGGTAGTCC

Standard DNA variants used in the fungal analysis (developed in this study)

>STD_fng1

CTTGGTCATTTAGAGGAAGTAAAAGTCGTAACAAGGCTTCCGTAGGTGAACCTGCGCAAGGATCATTAGGTATCACTCAGGAAGCAGACACAGAAAGACACGGTCTAGCAGATCGTTTATCGGCTAGGTCAAATAGAGTGCTTTGATATCAGCATGTCTAGCTTTAGAATTCAGTTTAGTGCGCTGATCTGAGTCGAGATAAAATCACCAGTACCCAAAACCAGGCGGGCTCGCCACGTACATCCAACAACGGATCTCTTGGTTCCGGCATCGATGAAGAACGCAGCGAA

>STD_fng2

CTTGGTCATTTAGAGGAAGTAAAAGTCGTAACAAGGATCCCGTAGGTGAACCTGCGTAAGGATCATTATCCTGCCCAGTAGCGGATGATAATGGTTGTTGCCAGCCGGTGTGGAAGGTAACAGCACCGGTGCGAGCCTAATGTGCCGTCTCCACCAACACAAGGCTATCCGGTCGTATAATAGGATTCCGCAATGGGGTTAGCAAATGGCAGCCTAAACGATATCGGGGACTTGCGATGACTTGCAACAACGGATCTCTAGGTTCCGGCATCGATGAAGAACGCAGCGAA

>STD_fng3

CTTGGTCATTTAGAGGAAGTAAAAGTCGTAACAAGGTATCCGTAGGTGAACCTGCGCAAGGATCATTAGAGGCGTTACCCCAATCGTTCAGCGTGGGATTTGCTACAACTTCTGAGTGCTACATGTACGAGACCATGTTATGTATGCACAAGGCCGACAATAGGACGTAGCCTTCGAGTTAGTACGTAGCGTGGTCGCATAAGCACAGTAGATCCTCCCCGCGCATCCTATTTATTAAGACGTTCAACAACGGATCTCTTGGTTCCGGCATCGATGAAGAACGCAGCGAA

>STD_fng4

CTTGGTCATTTAGAGGAAGTAAAAGTCGTAACAAGGCTACCGTAGGTGAACCTGCGTAAGGATCATTATTAATTCTATAGCAATACGATCATATGCGGATGGGCAGTGGCCGGTAGTCACACGTCTACCGCGGTGCTCAATGACCGGGACTAAAGAGGCGAAGATTATGGTGTGTGACCCGTTATGCTCGAGTTCGGTCAGAGCGTCATTGCGAGTAGTCGATTGCTTTCTCAATCTCCACCTACAACAACGGATCTCTGGGTTCCGGCATCGATGAAGAACGCAGCGAA

>STD_fng5

CTTGGTCATTTAGAGGAAGTAAAAGTCGTAACAAGGTTGCCGTAGGTGAACCTGCGCAAGGATCATTAGTACAGGTTCGCCTGTCGCCAAGATGCCTTACCTAGATGCAATGACGGACGTATTCCTCTGGCCTCAACGGTTCCTGCTTTCGCTGGGATCCAAGATTGGCAGCTGAAACCGCCTTTCCAAAGTGAGTCCTTCGTCTGTGACTAACTGTGCCAAATCGTCTTGCAAACTCCACATTCAACAACGGATCTCTCGGTTCCGGCATCGATGAAGAACGCAGCGAA
